# The lateral septum orchestrates state-dependent modulation of associative threat memory dynamics across the ovarian hormone cycle

**DOI:** 10.64898/2026.01.28.702128

**Authors:** Nina E. Baumgartner, Peter J. Teravskis, Neysa S. Dechachutinan, Kristen H. Adcock Binion, Gaven C. Bell, Amuktha S. Dasari, Gavin C. Newberry, Meghna Penumudi, Timothy H. Rumbell, Elizabeth K. Lucas

## Abstract

Cycling ovarian hormones orchestrate neural states that greatly impact brain function and guide behavior. Here, we discover that cued threat memory processes are regulated across the mouse estrous cycle in a state-dependent manner. We show that low hormone states lead to overexpression of threat memory in females. However, high hormone states confer protection against this overexpression via state-dependent recruitment and reengagement of the lateral septum (LS) to the memory ensemble. Chemogenetic manipulations confirmed the LS as necessary and sufficient to suppress female memory overexpression. Single nucleus sequencing revealed a novel LS neuron population, defined by coexpression of neurotensin and somatostatin, selectively recruited to the female memory ensemble during high hormone states. This estradiol-sensitive population exhibits a female bias in projection strength to the nucleus accumbens and displays unique calcium dynamics associated with state-dependent memory suppression. Altogether, we uncover a sexually divergent neural mechanism whereby cycling ovarian hormones modulate memory expression.

## INTRODUCTION

Women are twice as likely as men to experience post-traumatic stress disorder (PTSD)^1,2^. Clinical research demonstrates a strong influence of cycling ovarian hormones in driving this enhanced female susceptibility^3^. Menstrual cycle stage at the time of trauma influences the risk of subsequent PTSD development^4–6^, and symptom severity fluctuates in time with hormone levels across the menstrual cycle^7–9^. Thus, ovarian hormone states modulate the development and expression of fear-based psychiatric symptoms.

Fluctuating hormone levels across the human menstrual cycle or rodent estrous cycle orchestrate widespread changes in brain plasticity. Rising estradiol levels during the late follicular stage of the menstrual cycle drive temporary reorganization of functional brain networks, while the post-ovulatory progesterone surge opposes these effects^10,11^. Similar patterns of hormone fluctuations across the four-day rodent estrous cycle induce cyclic changes in neuronal morphology^12,13^, physiology^14,15^, and chromatin dynamics^16,17^. Crucially, these effects are not necessarily attributable to the actions of a single hormone^17,18^, but instead require the synchronization of various hormones across time. In rodents, surging ovarian estradiol levels during the day of proestrus stimulate subsequent secretion of hypothalamic^19^, pituitary^19^, adrenal^20^, and gastrointestinal^21–23^ hormones to drive changes in sexual^24^, appetitive^25^, and locomotor^26^ behaviors. How these fluctuating physiological states impact cognitive function remains hotly debated.

Cued threat conditioning is the most well-established preclinical model for investigation of the biological mechanisms underlying memory processes dysregulated in PTSD^27^. However, the few studies considering the rodent estrous cycle in this model have yielded inconsistent results^28^. While both genetic^29^ and environmental^30^ factors contribute to these inconsistencies, a major practical challenge of this work has been in the timing of experimental designs—deciding which aspect of the memory process to target to specific hormone states. We therefore approached this question through the framework of state-dependent memory, a long-observed phenomenon in which memory recall is most efficient when both memory acquisition and recall occur in the same physiological state^31^. This effect is attributed to shifts in neural network states that promote subcortical regulation of memory over cortical control^32^, an idea first proposed in 1937^31^ and convincingly demonstrated by more recent studies in mice^33,34^. Although classically associated with pharmacological manipulations, any endogenous or exogenous factors with widespread impacts on brain function and neuronal plasticity can produce physiological states that promote state-dependent memory^32^. Considering the dynamic changes in brain plasticity orchestrated by cycling ovarian hormones, we hypothesized that hormone fluctuations across the mouse estrous cycle would influence state-dependent memory processes.

Here, we investigate the behavioral and neural impacts of estrous cycle state on cued threat memory. We first establish that cued threat memory expression is regulated across the estrous cycle in a state-dependent manner. Low hormone states drive overexpression of cued threat memory, but high hormone states protect against this overexpression, rendering females experiencing memory acquisition and recall in proestrus behaviorally indistinguishable from males. We then use a powerful combination of modern neuroscience tools, including brain-wide quantification of neural activity, chemogenetics, single nucleus RNA sequencing (snSEQ), and calcium imaging, to uncover the mechanisms driving this effect. We discover that recruitment and reengagement of a novel neuronal population in the lateral septum (LS) limits overexpression of cued threat memory in females—but not males. Overall, this work uncovers a sexually divergent neural mechanism whereby the mouse estrous cycle modulates the fundamental biological processes of memory formation in a state-dependent manner.

## RESULTS

### Estrous cycle drives state-dependent modulation of cued threat memory in female mice

To test the hypothesis that ovarian hormone levels could influence state-dependent memory processes, we conditioned male mice and female mice in either high (proestrus, P) or low (diestrus, D) hormone states using auditory threat conditioning (Fig 1A). Mice received six co-terminating presentations of the conditioned stimulus (CS, 20 s tone) paired with the unconditioned stimulus (US, 2 s shock) spaced with 100 s inter-trial intervals (ITIs). We measured freezing, the primary defensive response displayed by rodents including C57Bl/6J mice of both sexes^29^, as the conditioned response^35,36^. All groups exhibited similar increases in CS-induced freezing over the course of threat conditioning (Fig 1B), with similar US reactivity levels (Fig S1A) and minimal CS-induced flight behaviors (Fig S1B). We then assessed cued recall, consisting of a 240 s baseline period followed by ten presentations of the CS with 100 s ITIs in a novel context, when female mice returned to the same (P→P or D→D) or opposite (P→D or D→P) estrous cycle stage. Freezing to the CS during recall was identical in male and P→P female mice (Fig 1C). However, any exposure to the low hormone state diestrus during either conditioning or recall resulted in increased CS-induced freezing during recall. In many instances, freezing bouts during recall were exceptionally long in these mice, with one continuous bout lasting the majority (≥10 s) of a single CS (Fig 1D). Importantly, this behavioral effect was not driven by the length of time between conditioning and recall (Fig 1E) or by group differences in conditioned flight behaviors (Fig S1C). Furthermore, all experimental groups exhibited limited pre-CS baseline freezing (Fig 1C) and similar freezing during the ITIs (Fig S1D), indicating that the observed differences in threat memory expression are specific to the CS and not due to a generalized increase in freezing. Finally, anxiety states have been shown to influence threat memory recall^37^, so we assessed avoidance behavior on the elevated plus maze two hours prior to recall (Fig S1E). While P→P females exhibited reduced avoidance behavior in the CS-only control group (Fig S1F-I), no group differences were observed in the conditioned group (Fig S1E), precluding any effects of anxiety states on freezing during cued recall.

**Figure 1.**
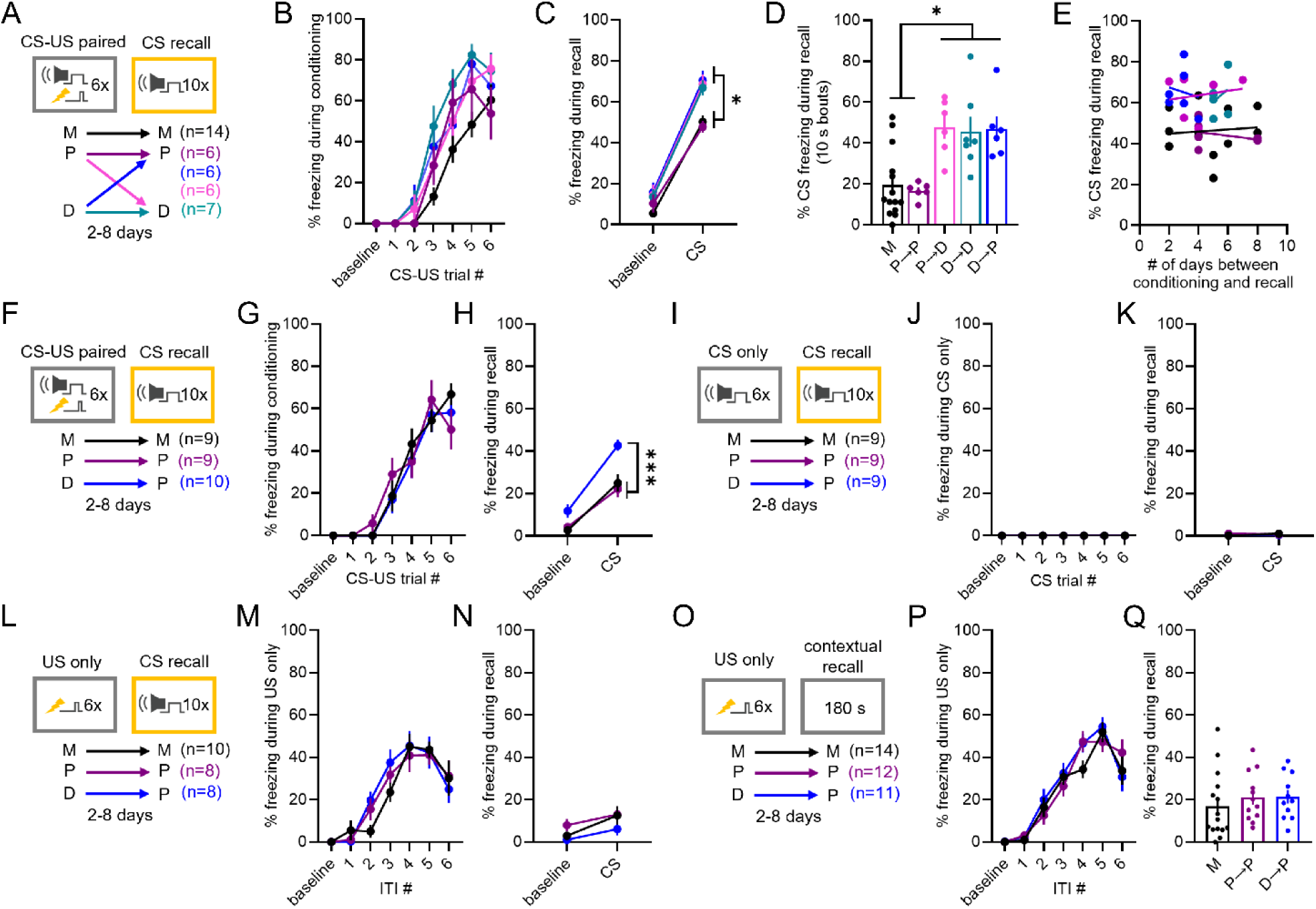
Estrous cycle drives state-dependent modulation of cued threat memory expression in female mice. **A-E)** The experimental schema is depicted in **(A)**. Male (M) mice or female mice in either high (proestrus, P) or low (diestrus, D) hormone states experienced auditory threat conditioning consisting of six co-terminating presentations of the conditioned stimulus (CS, 20 s tone) with the unconditioned stimulus (US, 2 s shock). Cued recall, consisting of ten CS presentations in a novel context, was tested when female mice returned to the same (P→P, D→D) or opposite (P→D, D→P) estrous stage. Percent of time freezing during the baseline and CS presentations across conditioning **(B)**, during the pre-CS baseline period and the average of all CS presentations during cued recall **(C)**, and for freezing bouts lasting ≥10 s during CS presentations in cued recall **(D)**. No significant correlations between freezing to the CS during recall and the length of time between conditioning and recall **(E)**. **F-H)** Following relocation of our laboratory, we repeated the prior CS-US paired experiment **(F)**. Percent of time freezing during the baseline and CS presentations across conditioning **(G)** and for the pre-CS baseline period and the average of all CS presentations during cued recall **(H)**. **I-K)** Experimental schema for CS-only control mice, which underwent identical experimental parameters as those in (F), except the US was omitted **(I).** Percent of time freezing during baseline and CS presentations across CS-only experience **(J)** and for the pre-CS baseline period and the average of all CS presentations during cued recall **(K)**. **L-N)** Experimental schema for US-only control mice, which underwent identical experimental parameters as those in (F), except the CS was omitted during conditioning **(L)**. Percent of time freezing during baseline and inter-trial intervals (ITIs) across US-only conditioning **(M)** and for the pre-CS baseline period and the average of all CS presentations during cued recall **(N)**. **O-Q)** To test for estrous cycle state-dependent effects of contextual threat memory, we performed context-forward conditioning and contextual recall consisting of 180 s in the conditioning context **(O)**. Percent of time freezing during baseline and ITIs across contextual conditioning **(P)** and during contextual recall **(Q)**. \**p* < 0.05, \*\*\**p* < 0.001 for Bonferroni-corrected post-hoc comparisons following a significant interaction effect in mixed-measures ANOVA **(C,H)** or significant one-way ANOVA **(D)**. For full statistics, see Table S1.

After completing this experiment, our laboratory moved to a different university. For replication and additional control experiments, we simplified the experimental groups to include males, P→P females, and D→P females so that female groups were in the same estrous stage at recall. Once again, D→P females exhibited enhanced CS freezing, whereas males and P→P females were behaviorally indistinguishable (Fig 1F-H). No group differences were observed in cued recall for CS-only (Fig 1I-K) and US-only (Fig 1L-N) control groups, eliminating effects of pseudoconditioning or sensitization. Finally, contrary to one study in rats^38^, we did not observe estrous cycle state-dependent regulation of contextual threat memory (Fig 1O-Q). Together, these experiments demonstrate that estrous cycle regulation of state-dependent memory is specific to associative learning processes. We posit that estrous cycle modulates the behavioral expression of associative threat memory, where a female bias towards exaggerated freezing responses is rescued during high hormone states.

**Figure S1.**
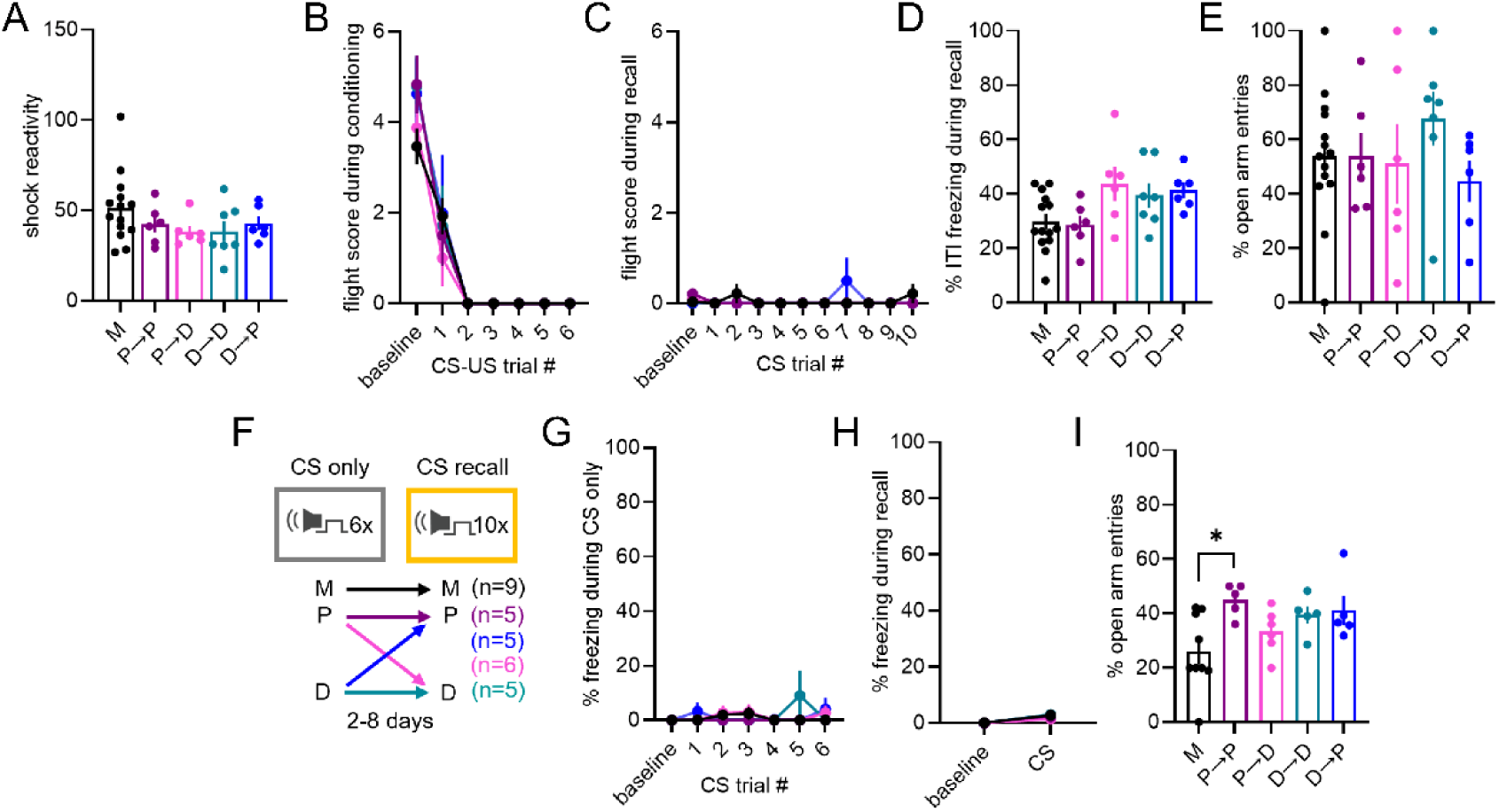
Related to Figure 1 – Additional behavioral control measures. **A-E)** Supporting data for Fig 1A-E. **A)** No group differences in shock reactivity to the US during conditioning. **B-C)** Minimal conditioned flight behaviors to the CS during conditioning **(B)** and recall **(C)**. **D)** No group differences in percent freezing during the ITIs at recall. **E)** Two hours prior to cued recall, mice were tested on the elevated plus maze, and no group differences in open arm avoidance were observed. **F)** Experimental schema. Prior to the relocation of our laboratory, we completed a CS-only control experiment identical to that in Fig 1A except the mice were never exposed to the US. **G-H)** CS-only controls exhibited minimal CS freezing during the CS-only experience **(G)** and minimal freezing during the pre-CS baseline period and the average of all CS presentations during cued recall **(H)**. **I)** Two hours prior to cued recall, mice were tested on the elevated plus maze. CS-only males exhibited fewer open arm entries compared to CS-only P→P females. \**p* < 0.05 for a Bonferroni-corrected post-hoc comparison following a significant one-way ANOVA. For full statistics, see Table S1.

### Identification of the LS as a state-dependent regional memory ensemble in high hormone states

Neuronal ensembles are engaged during learning and reactivated during recall, representing the physical trace of a memory in the brain^39^. Considering the profound influence of cycling reproductive hormones on neural activity^40,41^ and brain-wide functional states^11^, we hypothesized that state-dependent engagement of regional ensembles may alter memory expression across the estrous cycle (Fig 2A). As most of the canonical literature on associative threat memory circuitry has solely considered male brains, we chose an unbiased approach to assess estrous cycle regulation of regional brain activation following cued threat conditioning (Fig 2B) and recall (Fig 2C). Using protein expression of the neural activity marker c-fos as a proxy for neural activity, we obtained counts of c-fos+ cells from brain regions previously observed to be engaged by conditioning in both sexes^42^ in male, proestrus, or diestrus mice following cued threat conditioning or a naïve control experience (see Table S2 for full data). For any region displaying a significant estrous cycle-driven interaction post-conditioning, we also quantified c-fos expression following cued recall to identify state-dependent regional memory ensembles. Of the 114 regions analyzed post-conditioning, only 17 displayed significant interactions between conditioning and group. Of these interactions, only seven were driven by estrous cycle state (Fig 2D-J, Fig S2A-H). When we investigated the reactivation of these seven regions following cued recall (Fig 2D-J), only one displayed a significant interaction between conditioning and group: the rostral lateral septum (LSr, Fig 2E). We found that mice conditioned in proestrus displayed enhanced recruitment of the LSr compared to male and diestrus mice (Fig 2K, S2I). Remarkably, P→P females also displayed enhanced reengagement of the LSr during cued recall compared to all other groups (Fig 2L, S2J). While long known to inhibit maladaptive behavioral responses to imminent threats^43,44^, in recent years LSr has also been implicated in the regulation of behavioral responses to learned cued and contextual threats^45–50^. Therefore, we hypothesized that enhanced recruitment and reengagement of the LSr during high hormone states could coordinate the state-dependent protection against overexpression of threat memory in P→P females.

**Figure 2.**
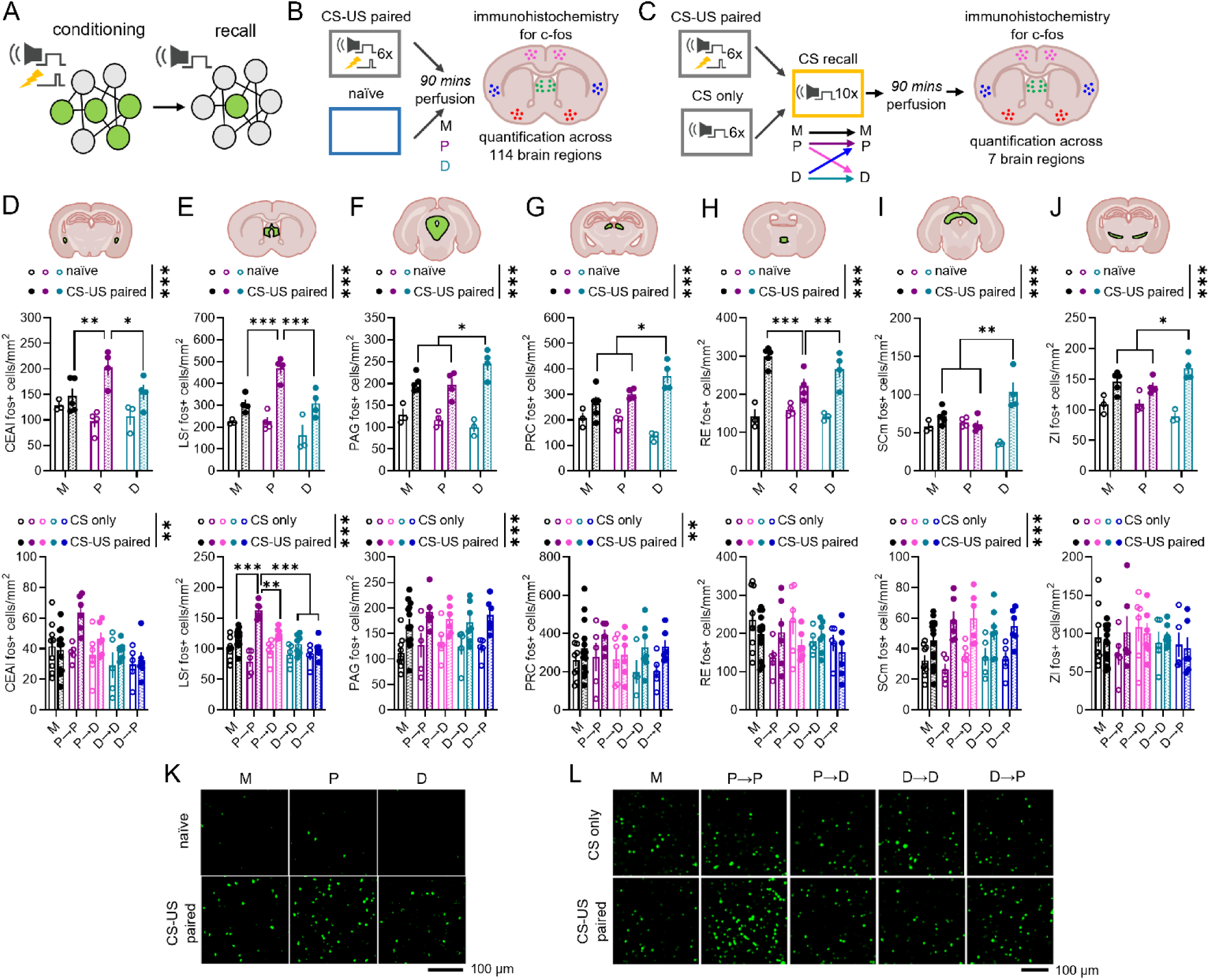
State-dependent recruitment and reengagement of the LSr to the female threat memory ensemble. **A)** Graphical depiction of the recruitment and reengagement of regional ensembles during threat conditioning and cued recall, respectively. **B)** Experimental schema of c-fos quantification post-conditioning. M, P, and D mice were randomly assigned to threat conditioned or naïve control groups. Ninety minutes post-conditioning, mice were perfused to perform immunohistochemistry and quantification of the neural activity marker c-fos across 114 brain regions outlined by the Allen Brain Atlas. **C)** Experimental schema of c-fos quantification post-recall. M, P, and D mice were randomly assigned to threat conditioned or CS-only control groups. When females were in the same or opposite estrous stage, mice were tested on cued recall and perfused ninety minutes later. **D-J)** c-fos counts of the seven regions showing significant group by conditioning interactions post-conditioning. Regional activation post-conditioning (top) and post-recall (bottom), for: **D)** lateral central amygdala (CEAl), **E)** rostral lateral septum (LSr), **F)** periaqueductal gray (PAG), **G)** precommissural nucleus (PRC), **H)** nucleus reuniens (RE), **I)** motor part of the superior colliculus (SCm), and **J)** zona inserta (ZI). Only LSr (E) showed a significant group by conditioning interaction post-recall. **K-L)** Representative images of LSr c-fos expression post-conditioning **(K)** and post-recall **(L)**. \**p* < 0.05, \*\**p* < 0.01, \*\*\**p* < 0.001 for main effects of conditioning (shown in legends above graphs) or for Bonferroni-corrected post-hoc tests following a significant interactive effect (shown on bar histogram) in a two-way ANOVA. For full statistics, see Table S2.

**Figure S2.**
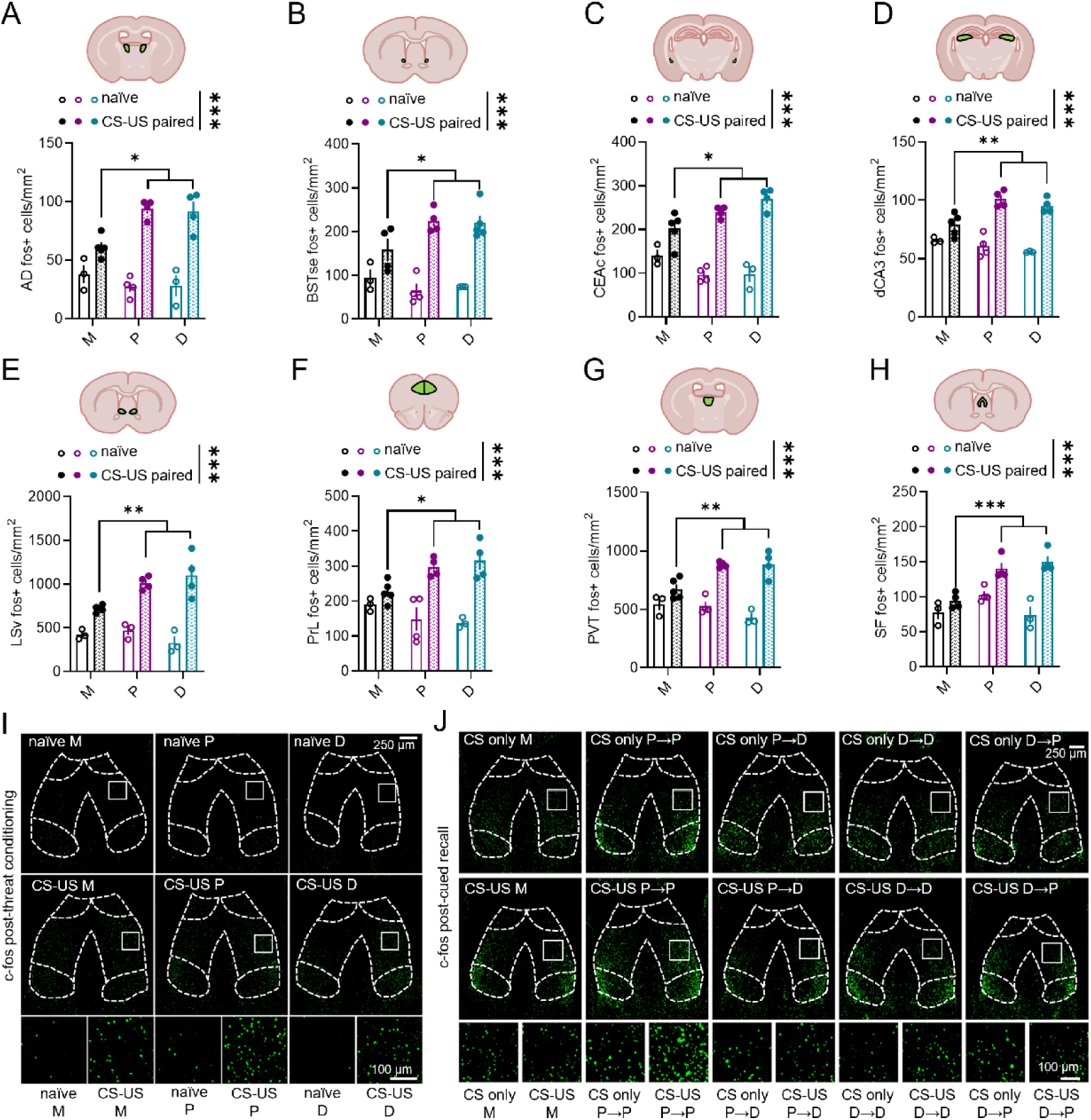
Related to Figure 2 - Additional regional c-fos analyses. **A-H)** c-fos expression in the eight brain regions displaying significant group by conditioning interactions post-threat conditioning that were driven by sex rather than estrous state, including: **A)** anterodorsal thalamic nucleus (AD), **B)** bed nucleus of the stria terminalis, strial extension (BSTse), **C)** capsular central amygdala (CEAc), **D)** dorsal CA3 of the hippocampus (dCA3), **E)** ventral lateral septum (LSv), **F)** prelimbic area (PrL), **G)** paraventricular thalamic nucleus (PVT), **H)** septofimbrial nucleus (SF). **I-J)** Representative images of c-fos expression across groups in the LS post-conditioning or naïve treatment **(I)** and post-cued recall in animals that previously underwent conditioning or a CS-only experience **(J)**. White boxes show the regions magnified in the insets below. \*\*\**p* < 0.001 for main effects of conditioning (shown in legends above graphs) or for Bonferroni-corrected post-hoc tests following a significant interactive effect (shown in bar histograms) in a two-way ANOVA. For full statistics, see Table S2.

In addition to analyzing regional memory ensembles, we also used the post-conditioning regional c-fos counts to conduct an exploratory analysis of changes in functional connectivity and neural network structure engaged by cued threat conditioning across groups^42^. We used covariance in regional c-fos expression within each group as a proxy for functional connectivity (Fig S3A) and applied graph theory derived analyses to assess neural network structures within each state (Fig S3B). Interestingly, the structure and composition of neural networks varied across groups, and unique hub regions were identified within each group (Fig S3C-D). Thus, mice experiencing the same behavioral experience under different hormone states exhibit distinct functional network structures following conditioning. Notably, LSr appeared as a hub region solely in the network of females conditioned in proestrus, bolstering our hypothesis that increased LSr activity may be key to the state-dependent suppression of threat memory overexpression in P→P females.

**Figure S3.**
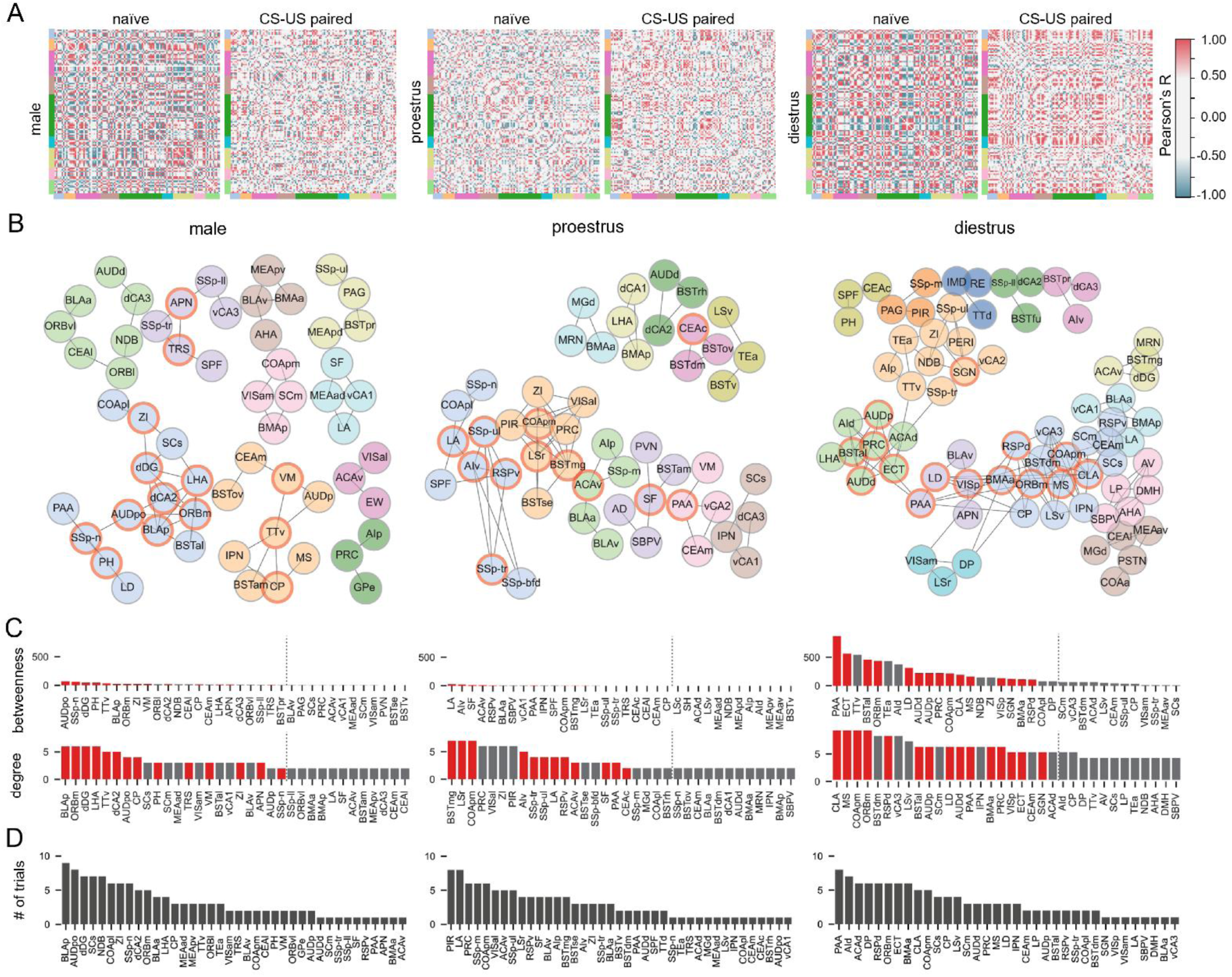
Related to Figure 2 – Exploratory functional connectivity and network analyses following threat conditioning. **A)** Correlation matrices of interregional c-fos expression across experimental groups (see schema in Fig 2B). Axes correspond to 114 brain regions organized by major divisions and color coded as indicated in Table S2. Matrix colors represent the correlation strength (Pearson’s R coefficients). **B)** Network graphs depicting brain regions (network nodes) as circles and supra-threshold correlations between regions as lines (network edges) for each CS-US paired group. A Markov clustering algorithm was applied to each adjacency matrix to identify clusters of nodes more connected with each other than with nodes outside the cluster. Node colors represent clusters. **C)** Bar histograms show the top 30% of nodes with the highest measures of betweenness (number of shortest paths between every pair of nodes in the network that traverse a node) and degree (number of edges to a node) for each network. Hub regions (depicted in red) of each network are nodes within the top 20% for both betweenness and degree (indicated by vertical dotted lines). **D)** Sequential *p*-value pruning revealed nodes most reliably identified as hub regions across 10 *p*-value thresholds (0.005 to 0.05) within each network.

### Enhanced LS activity is necessary and sufficient to promote state-dependent suppression of cued threat memory in females

As a major hub of the limbic system comprised primarily of inhibitory projection neurons^51^, the LS is well-suited to mediate the observed behavioral effects. Manipulations of LS activity in males have demonstrated a role for the region in both promoting^47–50^ and suppressing^45,46^ behavioral responses to learned cued or contextual threats. Our c-fos data (Fig 2E) suggest that enhanced recruitment and reengagement of LSr neurons may suppress behavioral responses to cued threats in females during high hormone states. Therefore, we decided to test the necessity and sufficiency of LSr activation in orchestrating the state-dependent modulation of threat memory across the estrous cycle.

We first hypothesized that inhibiting LSr activity during either conditioning or cued recall would result in increased freezing in P→P females. To test this hypothesis, we employed designer receptors exclusively activated by designer drugs (DREADDs) to chemogenetically inhibit LS activity (Fig 3A-B). We bilaterally injected an adeno-associated virus (AAV) expressing the inhibitory DREADD hM4D driven by a CaMKII promoter to target all projection neurons within the LS (Fig S4A). Thirty minutes prior to conditioning and/or recall, P→P female mice received an intraperitoneal injection of either the DREADD activator deschloroclozapine (DCZ) or vehicle (Fig 3C-E). There was no effect of DCZ on threat memory acquisition (Fig 3C). During recall, we observed a subtle effect of DCZ on pre-CS baseline freezing whereby mice receiving DCZ for both experiences exhibited increased freezing compared to mice receiving DCZ only during conditioning (Fig 3D). To control for this confound in our analysis of cued memory recall, we subtracted the percent of time freezing during the pre-CS baseline from the percent of time freezing to the CS for each mouse^52^. Remarkably, DCZ administration before conditioning and/or recall negated the state-dependent suppression of freezing in P→P females, resulting in increased CS freezing during recall (Fig 3E). When we repeated this experiment in males (Fig 3F-H), LS inhibition did not impact threat memory acquisition (Fig 3F) or CS freezing during recall (Fig 3H) but did induce a more robust state-dependent increase in pre-CS baseline freezing (Fig 3G) than observed in females, indicative of contextual generalization and consistent with previous work demonstrating an important role for the LS in contextual threat memory discrimination in males^46^. Importantly, expression and efficacy of DREADD-induced LS inhibition by DCZ was confirmed with c-fos immunostaining from mice perfused 90 minutes after cued recall and found to be equivalent between males and proestrus females (Fig 3I-K, S4B-C). Thus, we find that LS activity is necessary for the state-dependent suppression of freezing in P→P females but is not necessary for cued threat memory expression in males.

**Figure 3.**
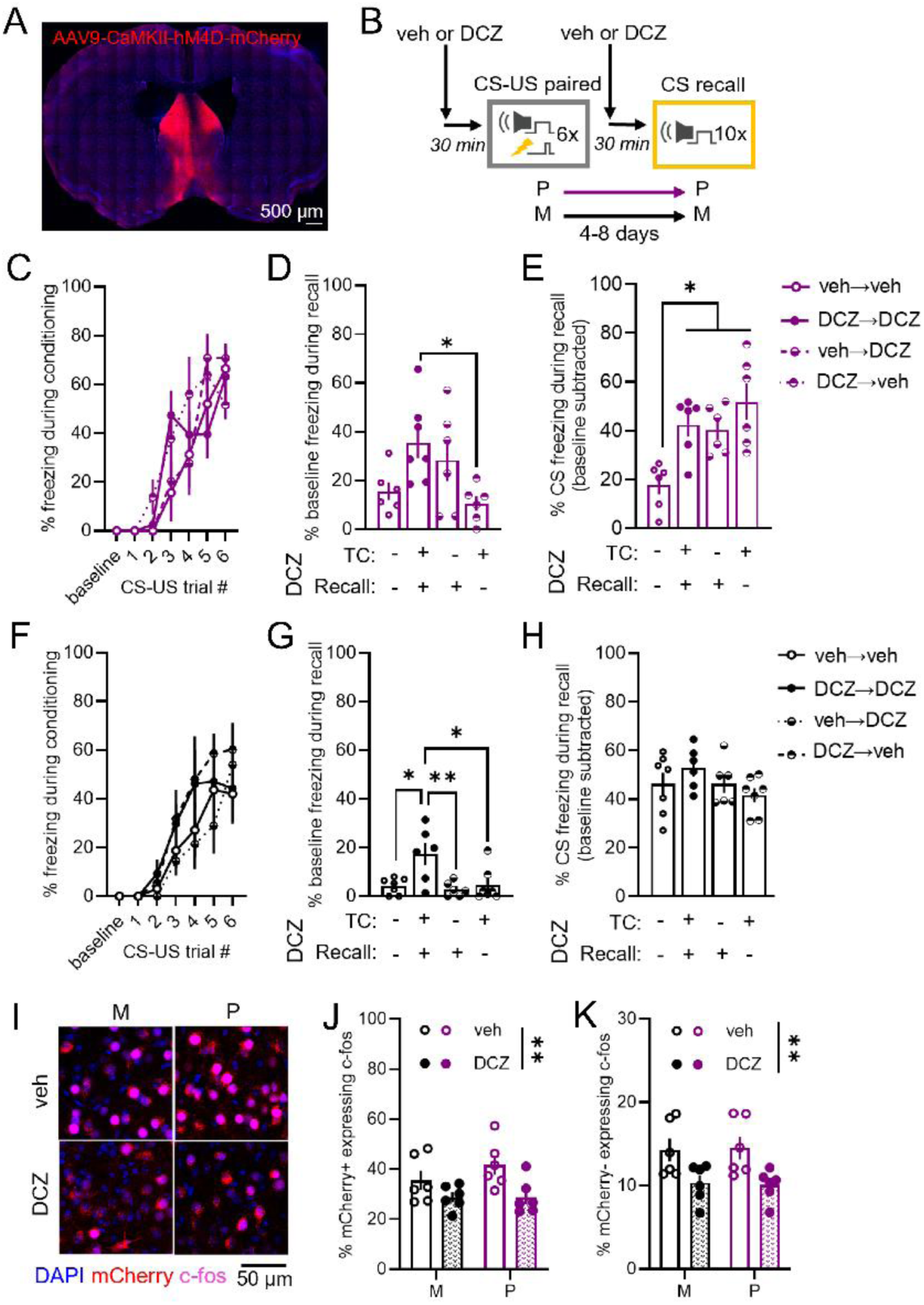
Activation of the LS is necessary for state-dependent inhibition of memory expression in high hormone states. **A-B)** Representative image **(A)** and experimental schema **(B)**. The inhibitory designer receptor exclusively activated by designer drugs (DREADD) hM4D was bilaterally expressed in LS projection neurons of females **(C-E)** and males **(F-H)**. Thirty minutes prior to threat conditioning and recall, P and M mice received intraperitoneal injections of either the DREADD activator deschloroclozapine (DCZ) or vehicle (veh). **C)** Percent of time freezing during baseline and CS presentations across conditioning for females. **D)** Percent of pre-CS baseline freezing during cued recall for females. Drug administration during threat conditioning (TC) and recall is denoted beneath bar histograms. **E)** Percent of time freezing to the CS during cued recall (with baseline freezing subtracted) for females. **F)** Percent of time freezing during baseline and CS presentations across conditioning for males. **G)** Percent of pre-CS baseline freezing during cued recall for males. **H)** Percent of time freezing to the CS during cued recall (with baseline freezing subtracted) for males. **I-K)** Ninety minutes following recall, mice were perfused and immunohistochemistry for c-fos was performed to validate DREADD efficacy. Administration of DCZ reduced the percentage of cells expressing c-fos in both mCherry+ **(J)** and mCherry- **(K)** cells. \**p* < 0.05, \*\**p* < 0.01 for Bonferroni-corrected post-hoc tests (shown in bar histograms) following a significant one-way ANOVA **(D,E,G)** or for main effects of treatment (shown in legends above graphs) following a significant two-way ANOVA **(J,K)**. For full statistics, see Table S3.

**Figure S4.**
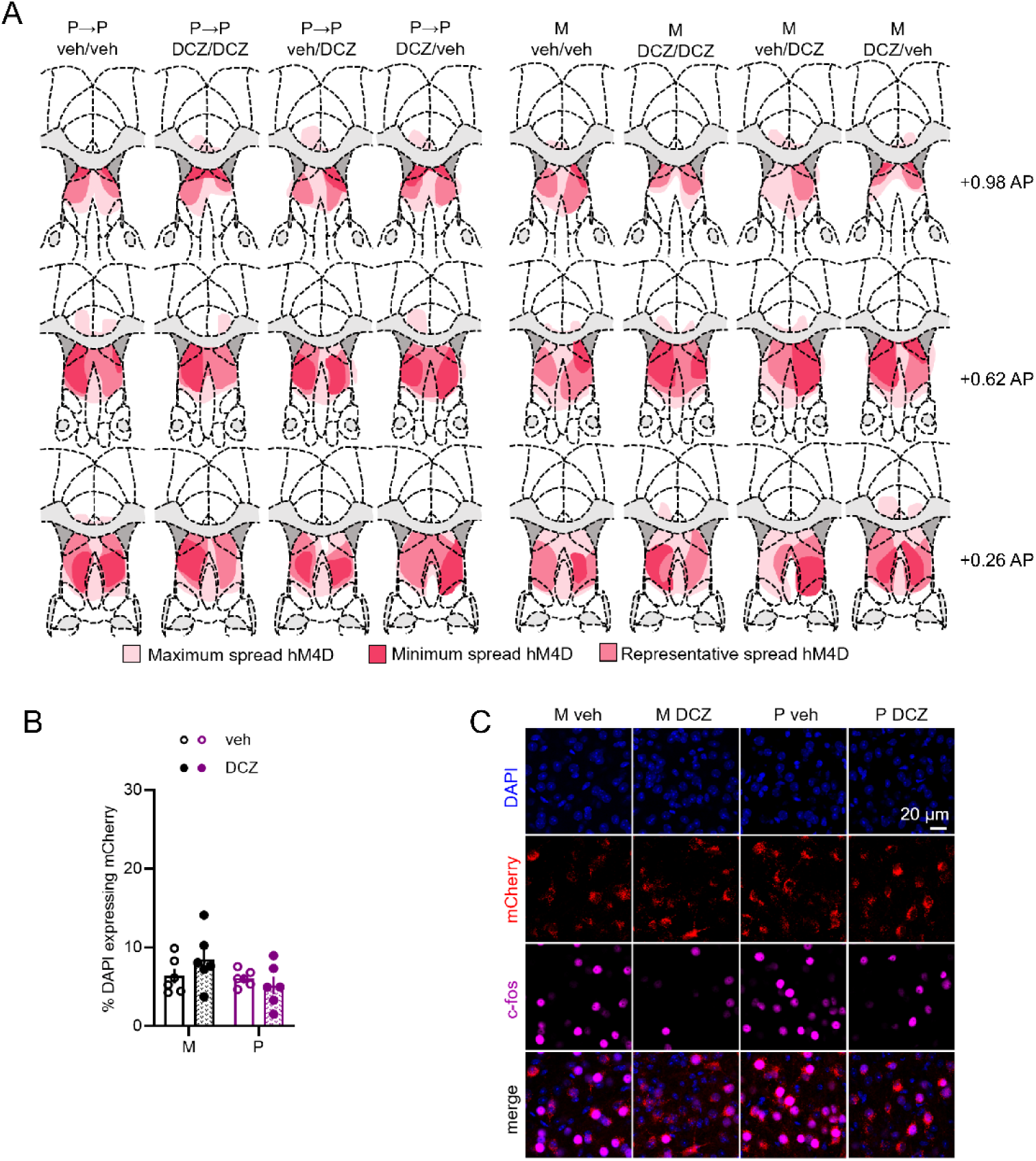
Related to Figure 3 - Supporting data for inhibitory DREADD experiments. **A)** Targeting maps showing hM4D-mCherry expression across all groups. Anterior-posterior (AP) coordinates relative to Bregma shown on right. **B-C)** DCZ or veh was administered 30 min prior to cued recall, and DREADD efficacy was validated by immunostaining for c-fos in mice perfused ninety minutes after cued recall. **B)** There were no group differences in mCherry expression relative to DAPI. **C)** Representative images of mCherry and c-fos expression in M and P mice with or without DCZ. For full statistics, see Table S3.

We next hypothesized that increasing LS activity during low hormone states would be sufficient to confer protection against the overexpression of cued threat memory in females. We employed a similar approach as above, this time using the excitatory DREADD hM3D with the activator DCZ administered during low hormone states (Fig 4A-B, Fig S5A). One hour prior to conditioning and recall, we administered either DCZ or vehicle via intraperitoneal injections. Again, we observed no group differences in threat memory acquisition among females (Fig 4C). During recall, we observed a subtle increase in pre-CS baseline freezing induced by DCZ in D→D females (Fig 4D) and therefore again subtracted baseline freezing from CS freezing. Activation of the LS during threat conditioning in D→P females or during recall in P→D females was sufficient to suppress CS freezing to similar levels as P→P controls (Fig 4E).

**Figure 4.**
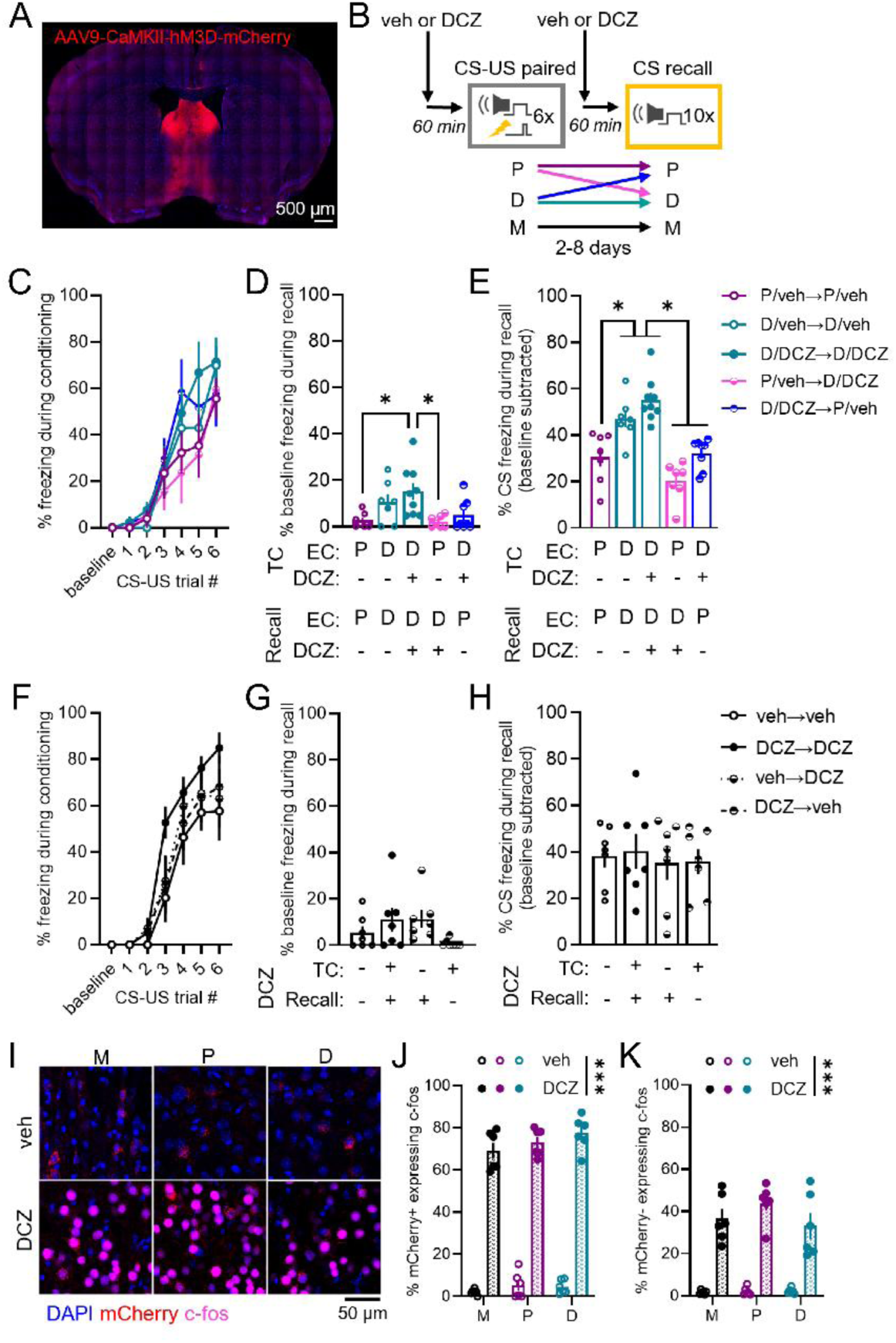
Activation of the LS is partially sufficient to suppress memory overexpression in low hormone states. **A-B)** Representative image **(A)** and experimental schema **(B)**. The excitatory DREADD hM3D was bilaterally expressed in LS projection neurons in females **(C-E)** and males **(F-H)**. One hour prior to TC and recall, P, D, and M mice received intraperitoneal injections of either the DREADD activator DCZ or veh. **C)** Percent of time freezing during baseline and CS presentations across conditioning for females. **D)** Percent of time freezing during the pre-CS baseline period of cued recall for females. Estrous cycle (EC) stage and drug administration during TC and recall are denoted beneath bar histograms. **E)** Percent of time freezing to the CS during cued recall (with baseline freezing subtracted) for females. **F)** Percent of time freezing during baseline and CS presentations across conditioning for males. **G)** Percent of pre-CS baseline freezing during cued recall for males. **H)** Percent of time freezing to the CS during cued recall (with baseline freezing subtracted) for males. **I-K)** Several days later, mice received another injection of either DCZ or veh followed by perfusion two hours later to validate DREADD efficacy with c-fos immunohistochemistry. Administration of DCZ increased the percentage of cells expressing c-fos in both mCherry+ **(J)** and mCherry- **(K)** cells. \**p* < 0.05, \*\*\**p* < 0.001 for Bonferroni-corrected post-hoc tests (shown in bar histograms) following a significant one-way ANOVA **(D,E)** or for main effects of treatment (shown in legends above graphs) following a significant two-way ANOVA **(J,K)**. For full statistics, see Table S4.

Surprisingly, LS activation was not sufficient to suppress freezing in D→D females, suggesting a distinct role for the physiological state of proestrus in mediating this behavioral effect beyond global LS activation. In males, LS activation via DCZ had no impact on threat memory acquisition (Fig 4F), pre-CS baseline freezing (Fig 4G), or CS freezing during recall (Fig 4H). Expression and efficacy of DREADD-induced LS activation by DCZ was confirmed following a separate injection of either DCZ or vehicle several days after behavioral testing and found to be equivalent across groups (Fig 4I-K, S5B-C). Thus, LS activity is sufficient to induce the state-dependent suppression of freezing in females experiencing either threat conditioning or cued recall in proestrus but has no effect on cued threat memory expression in males.

**Figure S5.**
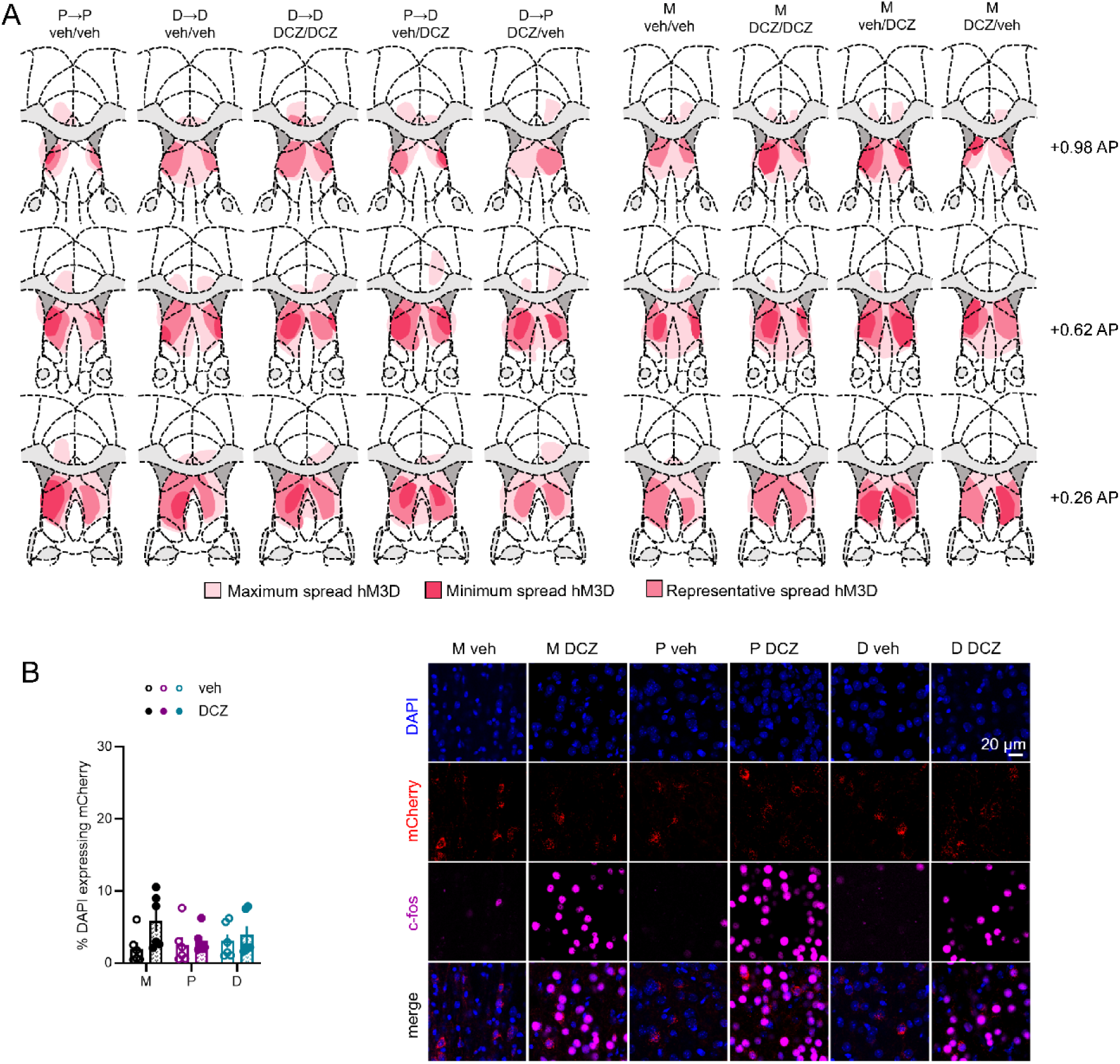
Related to Figure 4 - Supporting data for excitatory DREADD experiments. **A)** Targeting map showing hM3D-mCherry expression across all groups. AP coordinates relative to Bregma shown on right. **B-C)** DREADD efficacy was validated by immunostaining for c-fos in mice perfused two hours after intraperitoneal injections of either DCZ or veh several days following cessation of behavioral testing. **B)** There were no group differences in mCherry expression relative to DAPI. **C)** Representative images of mCherry and c-fos expression in M, P, and D mice with or without DCZ. For full statistics, see Table S4.

### LS*^Nts-Sst^* neurons are uniquely recruited to the female threat memory ensemble during high hormone states

As the LS is a highly heterogeneous region with a diverse range of cell types^53–57^, we posited that a specific neuronal population engaged during proestrus may be key to these state-dependent behavioral responses. In order to determine which populations are recruited to the threat memory ensemble, we performed snSEQ of the LS from proestrus females randomly assigned to either threat conditioned or naïve control groups. Five minutes after conditioning, micropunches of LS tissue were obtained and later processed for snSEQ (Fig 5A). After alignment and filtering in Cell Ranger, data sets underwent extensive pre-processing and quality control including background correction, removal of low-quality nuclei and doublets, and exclusion of off-target and splatter clusters (Fig S6A). Preliminary clustering revealed substantial technical variation across batches (clusters segregated by sample, poor integration between replicates, and differential expression of constitutive and marker genes between conditions) suggesting variability in transcript capture, cDNA amplification, and/or sequencing depth. To correct for batch-related artifacts, we applied a variational autoencoder model to separate the per-nucleus library-size scaling factor (𝓁) from biologically meaningful latent representations (𝔃) (Fig S6B)^58^.

**Figure 5.**
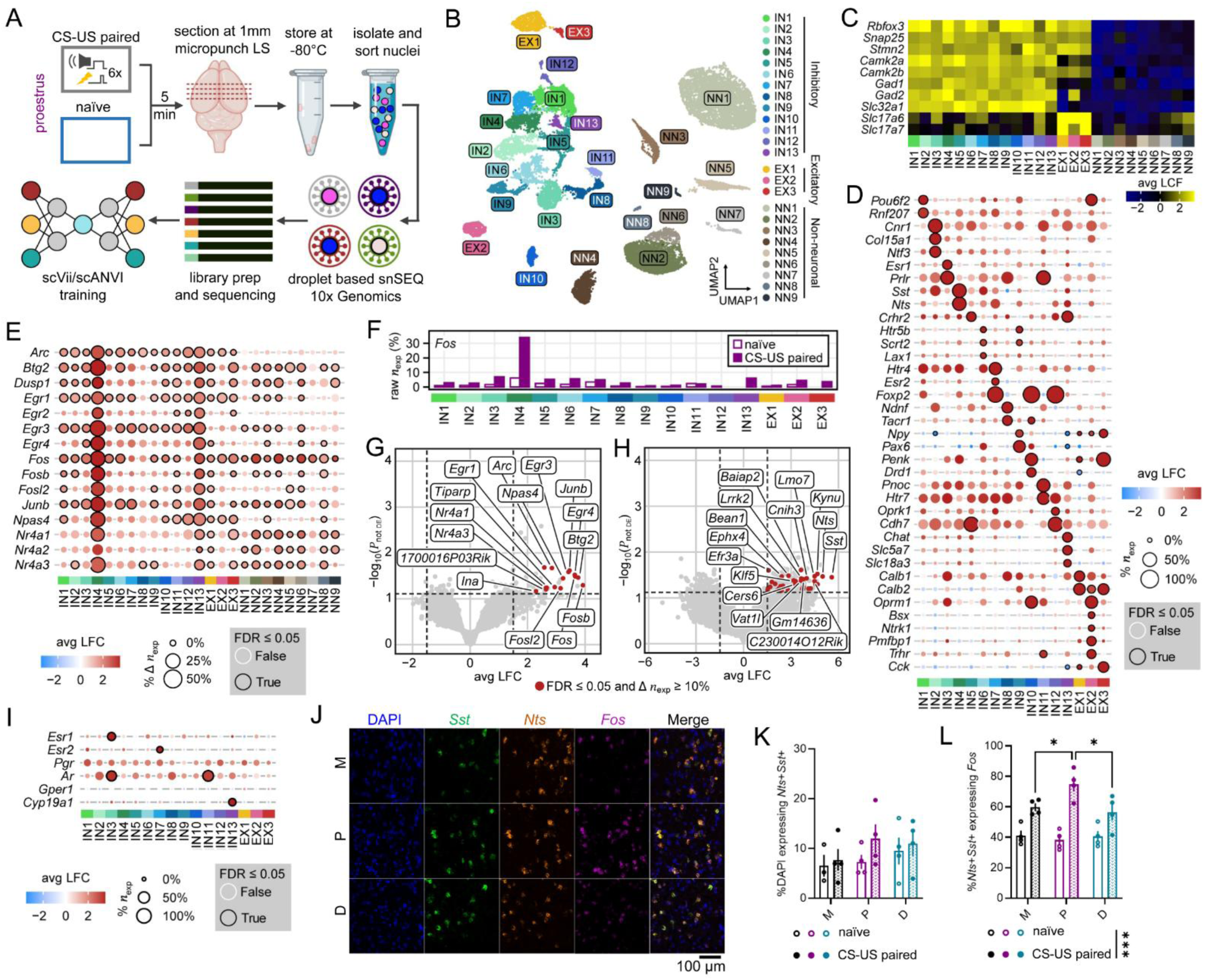
Recruitment of a novel inhibitory LS neuron population coexpressing neurotensin (Nts) and somatostatin (Sst) during threat conditioning in proestrus. **A)** Experimental schema. P mice were randomly assigned to threat conditioned or naïve control groups. Five minutes later, LS tissue was collected and later processed for single nucleus RNA sequencing (snSEQ) using 10x Genomics pipelines. For additional information on bioinformatics analyses, see Fig S6. **B)** Uniform manifold approximation and projection (UMAP) embedding of 25 cellular clusters spanning inhibitory neuron (IN), excitatory neuron (EX), and non-neuronal (NN) populations. **C)** Heatmap showing the average log_2_ fold change (avg LCF) of neuronal subtype marker expression across clusters. **D)** Dot plot of differential expression of selected cluster marker genes from cluster versus all other clusters comparisons. Dot colors indicate the avg LFC and dot sizes indicate the percentage of nuclei within each cluster that express each gene (%*n_exp_*). Genes meeting the false discovery rate threshold (FDR ≤ 0.05) are outlined in black. For full data, see Table S5. **E)** Dot plot of differential expression of selected immediate early genes from within-cluster comparisons. Dot sizes represent the percent of nuclei expressing each gene in the threat conditioned group relative to the naïve control group (%Δ *n_exp_*). For full data, see Table S6. **F)** Bar histogram of the percentage of nuclei in each cluster expressing at least one *Fos* transcript (raw *n_exp_ %*) in naïve and threat conditioned groups. **G-H)** Volcano plots of differential gene expression in IN4 for within-cluster naïve versus threat conditioned comparisons **(G)** and for cluster versus all other clusters comparisons **(H)**, with the top fifteen transcripts for each comparison labelled. Dashed vertical lines indicate a ±1.5 range from the avg LFC. Dashed horizontal lines indicate the FDR threshold. **I)** Dot plot of steroid hormone-related gene expression across neuronal clusters with dots demarcated as in **(D)**. **J-L)** Multiplex fluorescent *in situ* hybridization was performed on LS tissue from M, P, and D mice collected five minutes after threat conditioning or a naïve control experience. **J)** Representative images of LS from threat conditioned mice using probes for *Sst*, *Nts*, and *Fos*. **K)** Percentage of cells coexpressing *Nts* and *Sst*. **L)** Percentage of *Nts*+*Sst*+ cells expressing *Fos*. \**p* < 0.05, \*\*\**p* < 0.001 for Bonferroni-corrected post-hoc tests following a significant interactive (shown on bar histogram) or main effect (shown on legend) in a two-way ANOVA. For full *in situ* hybridization statistics, see Table S7.

**Figure S6.**
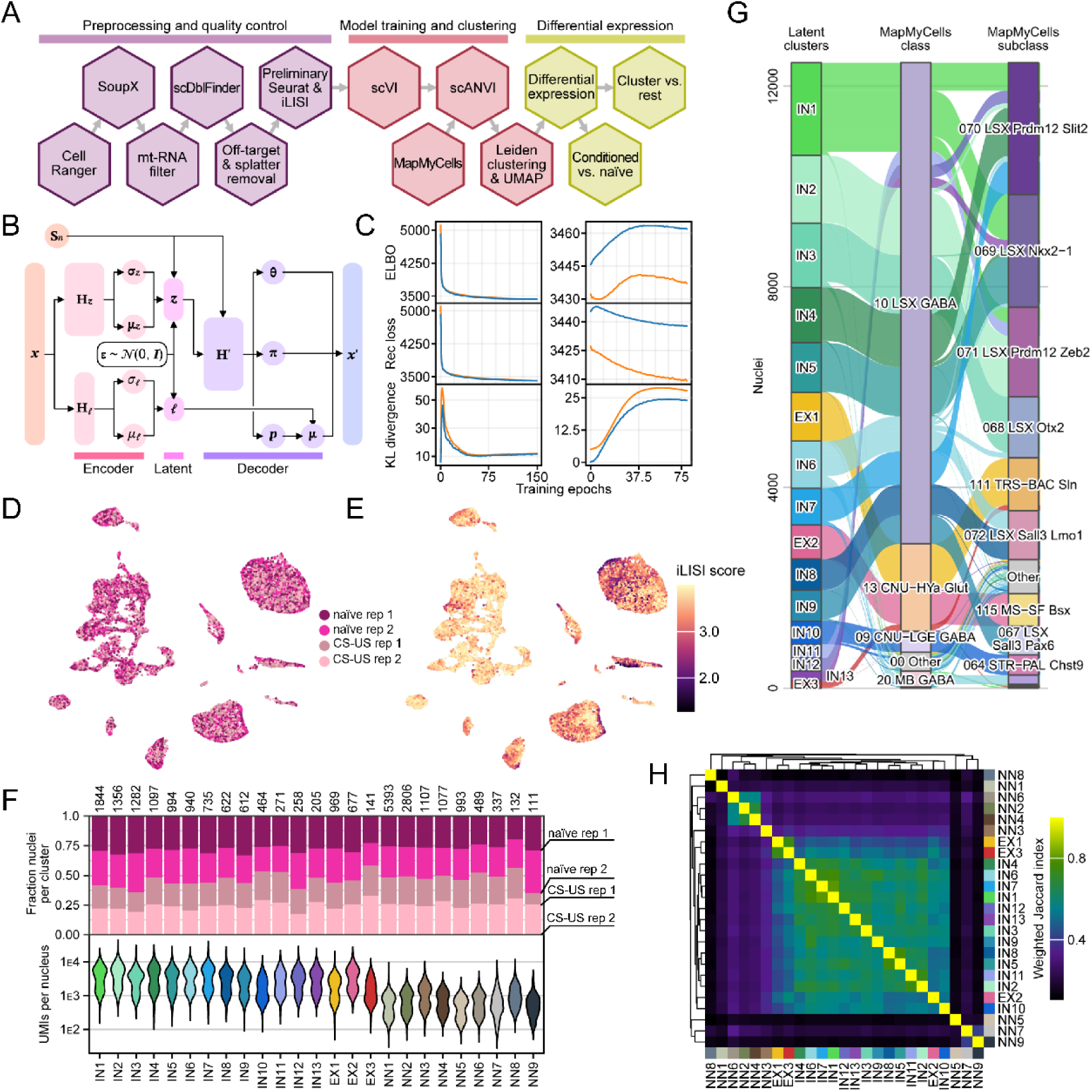
Related to Figure 5 – Supporting data for snSEQ analyses. **A)** Schematic of the snSEQ data processing pipeline. **B)** Simplified diagram of the single-cell variational inference (scVI) variable autoencoder (VAE) used for training, clustering, and differential expression analysis. Fine-tuning with single-cell annotation using variational inference (scANVI) extends this architecture by initializing an additional decoder module with extra output heads (not shown). **C)** Training and validation loss curves across epochs for scVI (left) and scANVI (right). Training loss is shown in blue, and validation loss in orange. **D)** UMAP of the scVI latent space colored by experimental condition and by replicate. **E)** UMAP of the scVI latent space colored by integration local inverse Simpson’s index (iLISI). The nearest-neighbor graph was computed from the z latent representation for each nucleus with k = 90. **F)** Stacked bar histograms showing the proportion of nuclei from each condition and replicate (rep) within each cluster, normalized to cluster size (top), and violin plots showing the distribution of total unique molecular identifiers (UMIs) per nucleus for each cluster (bottom). **G)** Sankey diagram linking neuronal clusters to their corresponding Allen Institute MapMyCells classes and subclasses **H)** Heatmap of weighted Jaccard indices for pairwise cluster-cluster comparisons of average fold change across all features derived from cluster versus all other clusters differential expression.

Following training and fine-tuning against Allen Brain Institute MapMyCells subtype annotations, Leiden clustering and uniform manifold approximation and projection (UMAP) on the latent space produced well-integrated results (Fig S6C-H). We identified 25 transcriptionally distinct clusters: 16 neuronal clusters expressing *Rbfox3* (13 inhibitory neuron clusters [IN1–13] expressing *Gad1*, *Gad2*, *Slc32a1*; 3 excitatory neuron clusters [EX1–3] expressing *Slc17a6* or *Slc17a7*) and 9 non-neuronal clusters identified by canonical glial, vascular, choroidal, and leptomeningeal markers (Fig 5B-D; Table S5). Comparison with prior LS snSEQ datasets^53–57^ confirmed that all neuronal clusters except IN6 corresponded to previously identified cell markers.

To determine which neuronal populations are recruited by conditioning in proestrus, we next performed differential expression analysis comparing a suite of immediate early genes (IEGs) in conditioned versus naïve nuclei across clusters. While modest IEG upregulation was observed across most clusters, two inhibitory clusters (IN4 and IN13) displayed markedly greater average expression and increases in the fraction of nuclei expressing IEGs (Fig 5E; Table S6). In particular, IN4 displayed a 5.52-fold increase in nuclei expressing *Fos* in conditioned versus naïve samples (Fig 5F). Notably, when comparing all differentially expressed genes in conditioned versus naïve samples, IN4 demonstrated broad IEG enrichment, including *Fosb*, *Btg2*, *Fos*, *Egr4*, *Junb*, *Egr3*, *Npas4*, *Fosl2*, *Arc*, *Egr1*, *Nr4a1*, and *Nr4a3* (Fig 5G), while IN13 showed selective enrichment of the IEGs *Egr4*, *Junb*, *Egr3*, *Npas4*, and *Arl5b* (Fig S7A). No other clusters displayed significant IEG enrichment within our analysis parameters (Table S6). The strongest markers for IN4 were somatostatin (*Sst*) and neurotensin (*Nts*; Fig 5H). LS*^Nts^* neurons and LS*^Sst^* neurons have separately been shown to suppress behaviors during stressful physiological states^57,59^ or in response to learned threats^45^, respectively. IN13 coexpressed corticotropin releasing hormone receptor 2 (*Crhr2*) along with inhibitory and cholinergic markers (*Chat*, *Slc5a7*, *Slc18a3*; Fig S7B). Surprisingly, neither of the clusters with IEG expression displayed enrichment in the steroid hormone receptors *Esr1*, *Esr2*, *Pgr*, *Ar*, or *Gper1* (Fig 5I); however, IN13 was enriched in *Cyp19a1* which encodes aromatase—the enzyme that converts testosterone to estradiol. In contrast to LS*^Nts^* and LS*^Sst^* neurons, previous studies have demonstrated that LS*^Crhr^*^2^ neurons promote defensive behaviors and increase physiological arousal^48,60^. Of note, we also identified a second population of *Crhr2*+ neurons (IN5) that did not exhibit IEG expression in response to conditioning (Fig S7C-D).

Therefore, we hypothesized that LS*^Nts-Sst^* neurons are selectively recruited to the proestrus memory ensemble. To test this hypothesis, we performed multiplex fluorescent *in situ* hybridization in male mice and female mice in proestrus or diestrus following threat conditioning or naïve control treatment. We analyzed *Fos* expression in LS*^Nts-Sst^* and LS*^Crhr^*^2^ neuron populations within the LSr (Fig 5J-L, S7E-G). No sex differences were observed for the proportion of LS cells from either population (Fig 5K, S7F). LS*^Crhr^*^2^ neurons exhibited widespread *Fos* expression independent of conditioning or group (Fig S7G). While the percentage of LS*^Nts-Sst^* cells expressing *Fos* was increased in all groups post-conditioning, proestrus females displayed enhanced LS*^Nts-Sst^* recruitment (Fig 5L), mirroring our pan-LSr c-fos cell counts (Fig 2E).

Therefore, we decided to target LS*^Nts-Sst^* neurons using an intersectional genetic approach to investigate the state-dependent recruitment of this population during conditioning and recall.

**Figure S7.**
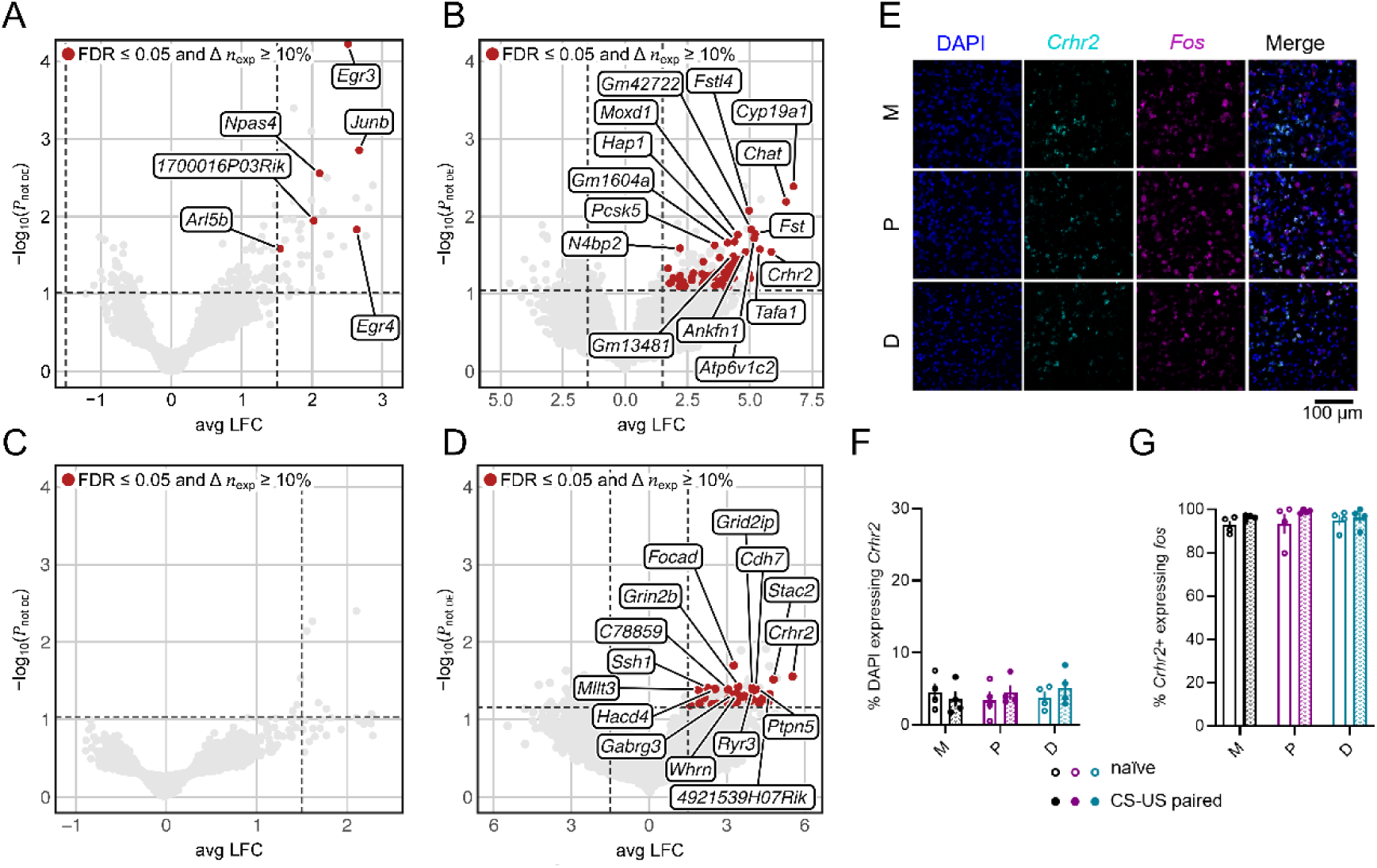
Related to Figure 5 – Additional data for corticotropin releasing hormone receptor 2 (Crhr2) expressing inhibitory neuron clusters. **A-B)** Volcano plots of differential gene expression for IN13, the other cluster showing IEG expression in Fig 5E, for within-cluster naïve versus threat conditioned comparisons **(A)** and for cluster versus all other clusters comparisons **(B)**, with the top fifteen transcripts for each comparison labelled when applicable. Dashed vertical lines indicate a ±1.5 range from the avg LFC. Dashed horizontal lines indicate the FDR threshold. **C-D)** Volcano plots of differential gene expression for the other *Crhr2*+ cluster (IN5) for within-cluster naïve versus threat conditioned comparisons **(C)** and cluster versus all other clusters comparisons **(D)**. **E)** Representative images of LS tissue from threat conditioned M, P, and D mice using fluorescent *in situ* probes for *Crhr2* and *Fos*. **F-G)** There were no group differences in the percentage of cells expressing *Crhr2* **(F)** or in the percentage of *Crhr2*+ cells expressing *Fos* **(G)**. For full statistics, see Table S7.

### State-dependent calcium dynamics of LS*^Nts-Sst^* neurons during threat memory processes

To better understand the temporal dynamics of LS*^Nts-Sst^* neuron recruitment to the threat memory ensemble, we generated double heterozygous *Nts*-Cre x *Sst*-FlpO mice to use for fiber photometry. Mice received unilateral LS infusions of an AAV conferring expression of GCaMP6f only in neurons expressing both Cre and Flp (COnFOn^61^) concurrent with fiber optic ferrule implants to measure bulk calcium transients from LS*^Nts-Sst^* neurons. Three weeks later, calcium activity was measured during conditioning and cued recall in males and P→P or D→P females (Fig 6A-B). Again, D→P females exhibited higher levels of CS freezing during recall, despite no differences in threat memory acquisition (Fig 6C). LS*^Nts-Sst^* neurons exhibited state-dependent patterns of recruitment across conditioning (Fig 6D-J). In P→P females, calcium responses were initially low and increased to both the CS (Fig 6E-F) and the US (Fig 6H-I) as the associative memory was formed. Conversely, in males and D→P females, responses to both stimuli were initially high and decreased over the course of conditioning. The rate of increased recruitment in P→P females during conditioning was predictive of behavior days later during recall, with higher degrees of recruitment to both the CS (Fig 6G) and the US (Fig 6J) associated with suppression of CS freezing during recall. During recall, P→P females also show heightened reengagement to the CS as compared to males and D→P females (Fig 6K-M).

**Figure 6.**
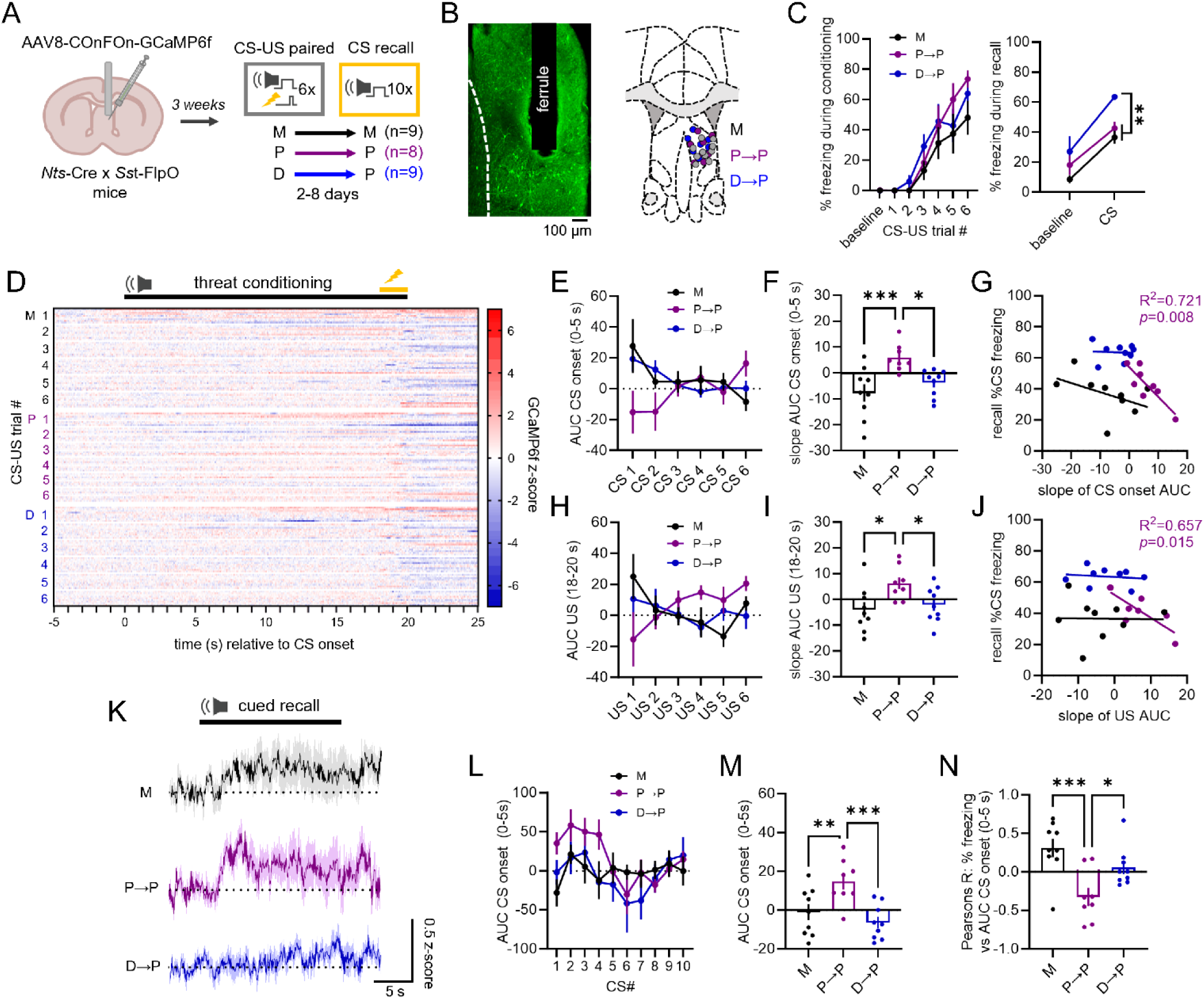
State-dependent calcium dynamics of LS^Nts-Sst^ neurons during threat memory processes. **A)** Experimental schema. Double heterozygous *Nts*-Cre x *Sst*-FlpO mice received unilateral LS injections of AAV8-COnFOn-GCaMP6f and optical ferrule implants three weeks prior to behavioral testing. Calcium activity of LS*^Nts^*^−^*^Sst^* neurons was recorded throughout conditioning and cued recall in M, P→P, and D→P mice. **B)** Representative image of viral expression and ferrule placement (left) and targeting map of ferrule placement across groups (right). **C)** Percent of time freezing to the CS during conditioning (left) and cued recall (right). **D)** Heatmap showing individual calcium responses across CS-US trials during conditioning. Black line represents time window of CS. Yellow line represents time window of US. Heatmap colors represent GCaMP6f z-scores. **E-G)** Area under the curve (AUC) for CS onset (0-5 s) was calculated across conditioning **(E)**, and the change in AUC across CS-US pairings was calculated as a slope **(F)**. The slope of AUC change was higher in P→P mice compared to M and D→P mice. **G)** In P→P mice alone, slope was negatively correlated with subsequent freezing to the CS days later during recall. Inset shows R^2^ and *p* values for linear regression of P→P within-subject correlation. **H-J)** AUC for US onset was similarly calculated across conditioning **(H)**, and P→P mice also displayed higher slopes of AUC change **(I)** that were negatively correlated with freezing to the CS during recall **(J)**. Inset shows R^2^ and *p* values for linear regression of P→P within-subject correlation. **K)** Average CS calcium responses during recall. Solid lines represent the group mean. Shading represents the SEM. Black line above traces represents the CS time window. **L)** AUC for CS onset across trials during cued recall. **M)** Average AUC for CS onset during cued recall. **N)** Within-subject correlations between calcium activity during CS onset and subsequent freezing for each CS presentation during recall. \**p* < 0.05, \*\**p* < 0.001, \*\*\**p* < 0.0001 for Bonferroni-corrected post-hoc comparisons following significant interactive effect in a mixed-measures ANOVA **(C)** or one-way ANOVA **(F,I,M,N)**. For full statistics, see Table S8.

Furthermore, within-subject correlations revealed that calcium responses to the CS during recall were negatively associated with freezing across individual trials in P→P females alone (Fig 6N). To verify the specificity of these patterns to LS*^Nts-Sst^* neurons, we repeated this experiment using a CaMKII-driven GCaMP6f to target all projection neurons within the LS (Fig S8A-C) and found minimal group differences during threat conditioning (Fig S8D-J) or recall (Fig S8K-N). Thus, LS*^Nts-Sst^* neurons display distinct recruitment patterns during threat memory acquisition in a hormone state-dependent manner, and both the degree of that recruitment and subsequent reengagement during recall is negatively associated with CS freezing during recall.

**Figure S8.**
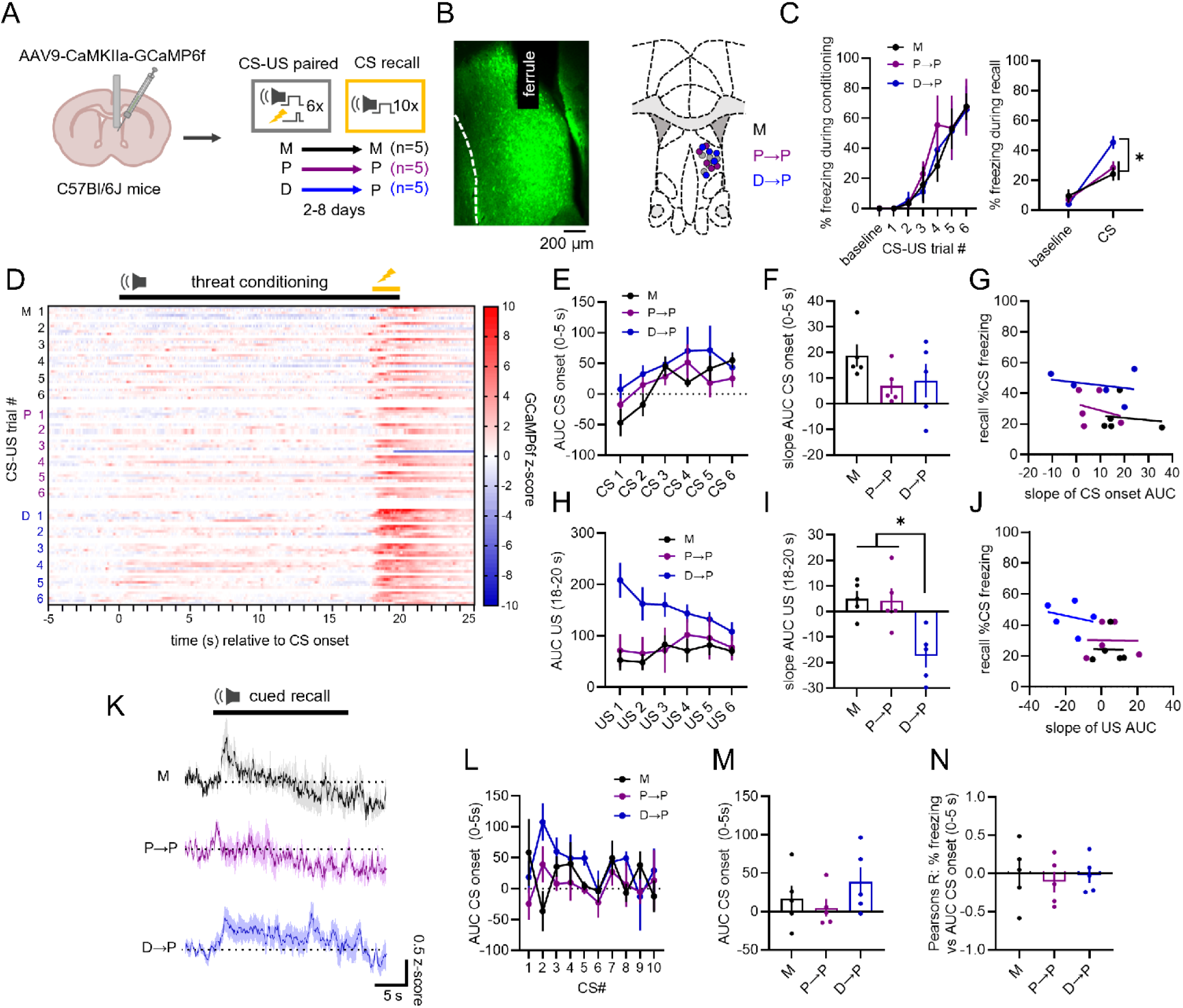
Related to Figure 6 – Global LS calcium dynamics during threat memory processes. **A)** Experimental schema. Mice received unilateral injections of AAV9-CaMKIIa-GCaMP6f and optical ferrule implants directed to the LS three weeks prior to behavioral testing. Calcium activity of LS projection neurons was recorded throughout conditioning and cued recall in M, P→P, and D→P mice. **B)** Representative image of viral expression and ferrule placement (left) and targeting map of ferrule placement across groups (right). **C)** Percent of time freezing to the CS during conditioning (left) or cued recall (right). **D)** Heatmap showing individual calcium responses across CS-US trials during conditioning. Black line above represents time window of CS. Yellow line represents time window of US. Heatmap colors represent GCaMP6f z-scores. **E-G)** AUC for the CS onset (0-5 s) was calculated across conditioning **(E)**, and the change in AUC across CS-US pairings was calculated as a slope. There were no group differences in slope of AUC to CS onset **(F)**, and no significant correlations of slope with subsequent freezing during cued recall **(G)**. **H-J)** AUC for the US onset was similarly calculated across conditioning for each group **(H)**. D→P female mice had significantly lower slope of AUC to US onset compared to M and P→P female mice **(I)**. However, there were no significant correlations of slope with subsequent freezing during cued recall **(J)**. **K)** Average CS calcium responses during recall. Solid lines represent the group mean. Shading represents the SEM. Black line above traces represents time window of CS. **L)** AUC for CS onset across trials during cued recall. **M)** Average AUC for CS onset during cued recall. **N)** Within-subject correlations between calcium activity during CS onset and subsequent freezing for each CS presentation during recall. \**p* < 0.05 for Bonferroni-corrected post-hoc comparisons following significant interactive effect in a mixed-measures ANOVA **(C)** or one-way ANOVA **(I)**. For full statistics, see Table S8.

### Characterization of LS*^Nts-Sst^* circuitry and recruitment

To better understand how LS*^Nts-Sst^* neurons guide threat memory expression, we next investigated their downstream projection targets. Double heterozygous male and female *Nts*-Cre x *Sst*-FlpO mice received bilateral LS infusions of a COnFOn-mCherry reporter and were perfused six weeks later to quantify mCherry terminals throughout the brain (Fig 7A-B). Unsurprisingly, we observed mCherry terminals in many brain regions involved in limbic function (Fig 7C-D). Consistent with previous investigations into LS*^Nts^* ^57,59,62,63^ and LS*^Sst^* ^45,64^ populations separately, we found LS*^Nts-Sst^* neurons project to the lateral hypothalamus, bed nucleus of the stria terminalis, and periaqueductal gray, among others. Therefore, LS*^Nts-Sst^* neurons are positioned to exert influence over limbic system function and ultimately regulate behavior through a variety of efferent connections.

**Figure 7.**
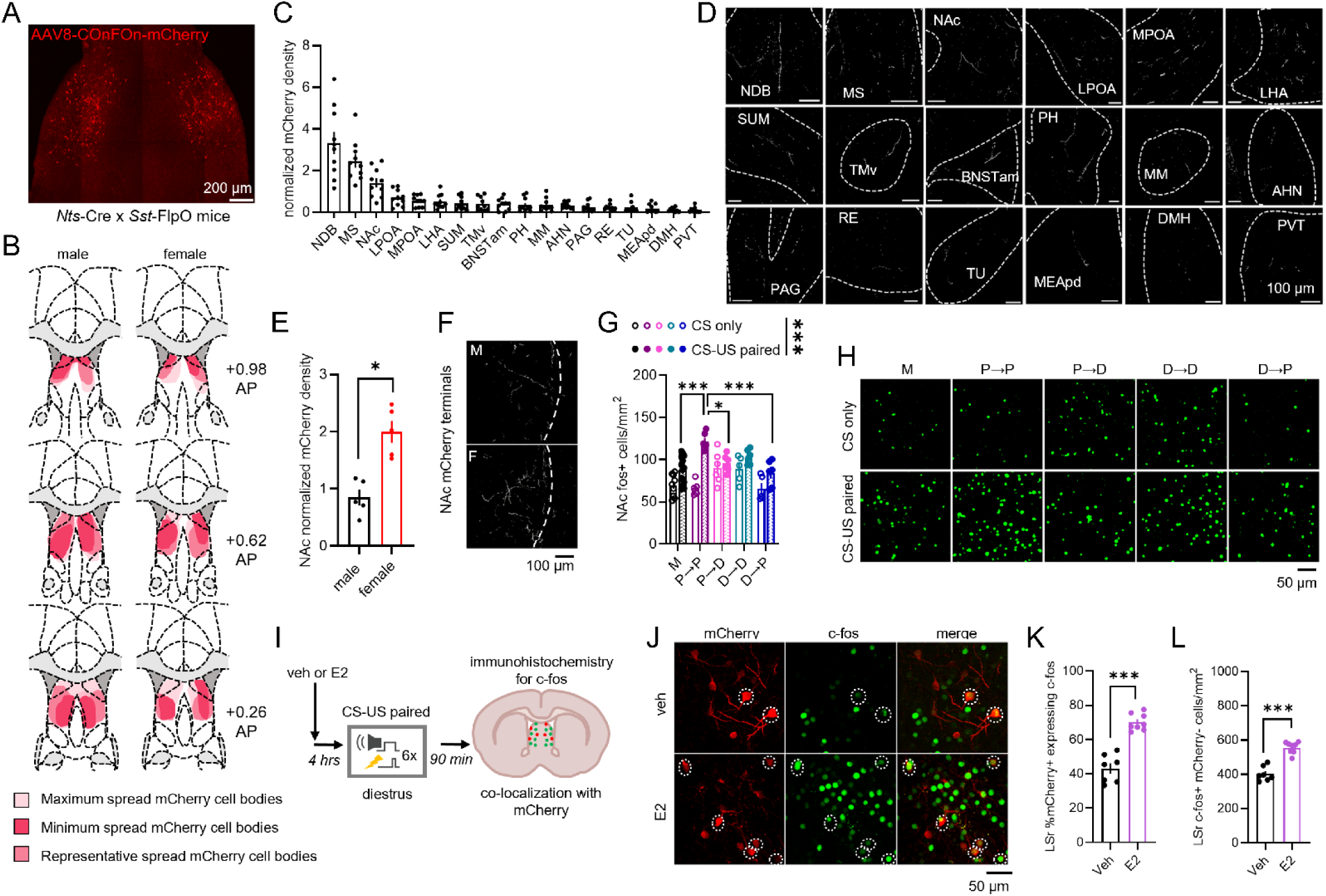
Characterization of LS^Nts-Sst^ circuitry and recruitment. **A)** Male and female double heterozygous *Nts*-Cre x *Sst*-FlpO mice received bilateral injections of AAV8-COnFOn-mCherry into the LS. **B)** Targeting map of somatic mCherry expression in each group. **C-D)** Quantification **(C)** and representative images **(D)** of mCherry terminals in brain regions in which at least four mice exhibited mCherry terminal expression. Density of terminals was normalized to area and the number of LS mCherry somas. Scale bars represent 100 μm for each image. **E-F)** Only the nucleus accumbens (NAc) displayed a sex difference in projection strength that survived Bonferroni correction **(E)**. Representative images of male (top) and female (bottom) NAc mCherry expression **(F)**. **G-H)** Quantification of regional c-fos expression post-recall from a prior experiment (see Fig 2C) revealed increased c-fos in the NAc of P→P females **(G)**. Representative images of c-fos expression **(H)**. **I)** Experimental schema. Female *Nts*-Cre x *Sst*-FlpO mice received bilateral injections of AAV8-COnFOn-mCherry into the LS. Three weeks later, upon entering diestrus, mice received subcutaneous injections of estradiol (E2) or vehicle (veh) four hours prior to threat conditioning. Mice were perfused ninety minutes post-conditioning, and immunohistochemistry was performed to quantify c-fos coexpression with mCherry. **J)** Representative images of mCherry and c-fos coexpression. Examples of mCherry+ neurons expressing c-fos are circled in white. **K-L)** E2 treatment prior to conditioning increased recruitment of both mCherry+ **(K)** and mCherry- **(L)** cells in the LSr. \**p* < 0.05, \*\*\**p* < 0.001 for Bonferroni-corrected *t*-tests (E), two-tailed *t*-tests **(K,L)**, or main effect of conditioning (shown in legend above graph) or Bonferroni-corrected post-hoc comparisons following a significant interactive effect (shown in bar histogram) in a mixed-measures ANOVA **(G)**. For full statistics, see Table S9.

Interestingly, we observed a robust sex difference in LS*^Nts-Sst^* projection strength to the nucleus accumbens (NAc), whereby females exhibited double the mCherry terminal density of males (Fig 7E-F; for all sex comparisons, see Table S9). This finding prompted us to quantify NAc c-fos expression in the brains of our cued recall experiment (Fig 2C). Remarkably, we detected increased NAc activation in P→P females (Fig 7G-H), paralleling what we observed in the LSr (Fig 2E) and suggesting a potential functional impact of this sex difference in LS*^Nts-Sst^* circuitry.

Lastly, we sought to uncover potential mechanisms driving enhanced recruitment of LS*^Nts-Sst^* neurons. While LS*^Nts-Sst^* neurons do not express estrogen receptors (Fig 5I), rising levels of estradiol initiate subsequent surges of hypothalamic, pituitary, adrenal, and gastrointestinal hormones that are also upregulated during proestrus. Thus, we hypothesized that administration of estradiol prior to threat conditioning would increase recruitment of LS*^Nts-Sst^* neurons in diestrus females through indirect estradiol action. Using colocalization of the COnFOn-mCherry reporter with immunostaining for c-fos (Fig 7I), we found that a subcutaneous injection of estradiol four hours prior to conditioning was sufficient to increase activation of LS*^Nts-Sst^* neurons (Fig 7J-K) and overall LSr activity (Fig 7L) in D females. Thus, the surge of estradiol during proestrus facilitates recruitment of LS*^Nts-Sst^* neurons to the female threat memory ensemble.

## DISCUSSION

Here, we employed the framework of state-dependent learning to demonstrate for the first time that fluctuating hormones across the mouse estrous cycle orchestrate brain states to modulate associative threat memory and guide behavioral responses to learned threats. Across numerous experimental designs and manipulations, we show that low hormone states bias females towards overexpression of cued threat memory but that high hormone states confer protection against this overexpression in a state-dependent manner, rendering females experiencing threat memory acquisition and recall in proestrus behaviorally indistinguishable from males. Unbiased assessment of neural activity unveiled the LSr as a regional memory ensemble when both memory acquisition and recall occur during high hormone states. Using bidirectional chemogenetic manipulations, we demonstrate that the LS is both necessary and sufficient for modulation of threat memory expression across the ovarian hormone cycle in females—with limited behavioral effects in males. Finally, we identify a novel estradiol-sensitive neuronal population in the LS that exhibits a female bias in projection strength to the NAc. Defined by coexpression of *Nts* and *Sst*, the activity of LS*^Nts-Sst^* neurons is associated with state-dependent suppression of threat memory expression in proestrus females. We posit that the heightened importance of the LS in female threat memory dynamics represents a “convergent” sex difference whereby LS engagement during high hormones states compensates for an otherwise “divergent” sex difference in behavior during low hormone states^65^.

Intriguingly, despite robust behavioral differences, very few brain regions exhibited state-dependent recruitment following conditioning, and only the LSr exhibited state-dependent reengagement following recall. A major center of the limbic system, the LS is known to integrate cognitive and affective information to drive appropriate behavioral and physiological responses to diverse stimuli^51^. Prior studies have reported increased neuronal excitability in the LS during high hormone states^66,67^, which may in turn prime recruitment of this region to the memory trace^68^. Consistent with the theory that state-dependent learning involves a shift in network dynamics away from cortical regulation towards subcortical control^32^, our neural network analyses identified the LSr as a proestrus-specific memory hub region that may drive such shifts in neural network structure. Interestingly, chemogenetic manipulations of LS projection neurons did not impact cued recall in males, contrary to prior studies that show LS inhibition can either increase^49^ or decrease^50^ freezing. However, LS inhibition did elicit a state-dependent increase in contextual threat generalization, indicating that state-dependent learning may contribute to the previously reported LS regulation of contextual threat memory discrimination in males^45–47^. In contrast to our findings in males, we report robust bidirectional effects of chemogenetic LS manipulations on cued threat memory in females whereby LS activation is both necessary and partially sufficient to induce state-dependent threat memory expression across the estrous cycle. Thus, enhanced LS activity in females is required to induce the same behavioral response to learned threats as males.

The neuronal composition of the LS is extremely heterogeneous^53–57^, so we performed snSEQ for unbiased identification of neurons recruited to the proestrus memory ensemble. Differential gene expression of a panel of IEGs revealed only two neuronal populations activated in response to threat conditioning. One is a previously identified population (LS*^Crhr^*^2^) known to promote autonomic and behavioral responses to threats^48,60^. The second population expresses both *Nts* and *Sst* (LS*^Nts-Sst^*), likely representing a subset of each respective population that has been previously characterized. LS*^Sst^*neurons are responsive to diverse aversive stimuli^45,63,64^ and suppress context-independent freezing in males^45^. LS*^Nts^* neurons are hyperactive during stressful physiological states^57,59,63^ and promote active, but not passive, behavioral responses to imminent threats^59^. Interestingly, LS*^Nts^* neurons were recently hypothesized to drive adaptive behavioral responses when an animal is in a weakened or stressful state^57^. Consistent with these findings, our *in situ* hybridization data confirmed that the LS*^Nts-Sst^*, but not the LS*^Crhr^*^2^, population is uniquely engaged by threat conditioning in proestrus.

Employing an intersectional genetic approach with *Nts*-Cre x *Sst*-FlpO mice, we made several important discoveries about LS*^Nts-Sst^* neurons. First, distinct patterns of recruitment and reengagement of LS*^Nts-Sst^* neurons during proestrus provide a neural correlate for the state-dependent suppression of cued threat memory in females. Second, administration of estradiol prior to threat conditioning is sufficient to enhance recruitment of these neurons during diestrus. Although our sequencing data do not show enrichment of ovarian hormone receptors in this population, LS*^Nts-Sst^* activity may be modulated by any of the numerous signaling pathways upregulated by estradiol during proestrus including various hormones^19–23^, neurotransmitters^69^, endocannabinoids^70^, and neurotrophins^71^. On the other hand, estradiol is known to enhance excitatory drive in the hippocampus^72,73^, which densely expresses estrogen receptors^74,75^ and constitutes the majority of synaptic inputs to the LS^76^. Thus, enhanced excitatory drive from upstream brain regions such as the hippocampus may also facilitate LS*^Nts-Sst^* recruitment to the memory ensemble under high hormone states. Finally, we found that LS*^Nts-Sst^* neurons are positioned to influence limbic system function through a broad range of efferent projections. Interestingly, we observed strengthened female projections to the NAc, which is strongly implicated in active avoidance learning^77,78^ and has been proposed as a “toggle switch” between passive and active responses to threats^79^. As we found state-dependent increases in NAc c-fos expression following recall in the same group exhibiting suppressed freezing and higher LS activity, this LS*^Nts-Sst^→*NAc circuit is well-poised to drive flexibility in behavioral responses to learned threats across hormone states.

A fundamental unanswered question is why threat memory expression would exhibit such dynamic modulation across the ovarian hormone cycle. Sometimes called “behavioral estrus”^80^, the high hormone state of proestrus drives physiological and behavioral changes, such as decreased avoidance^81,82^ and increased reward seeking^83,84^, that are necessary to find a mate and maximize reproductive success during ovulation. Similarly, while overexpression of threat memory may enhance survival during low hormones states when mating will not be fruitful, the suppression of threat memory during proestrus likely enhances reproductive fitness. Thus, selection pressure represents one explanation for the emergence of state-dependent threat memory expression via engagement of the female-specific LS memory ensemble. These findings highlight the dynamic nature of sex differences in neural function, whereby sexually divergent neural mechanisms are required to account for the behavioral flexibility exhibited by females across the ovarian hormone cycle.

In conclusion, we report an endogenous neural mechanism that suppresses the behavioral expression of threat memory in females. These findings highlight the complex influence of cycling reproductive hormones on neural function and illustrate the benefit of embracing, rather than discounting, sex-related biological variables in neuroscience research. Given the clinical evidence implicating ovarian hormone states in modulating the development and expression of PTSD, further investigation into this state-dependent neural mechanism may provide novel biomarkers and therapeutic targets that can be harnessed for the diagnosis and treatment of psychiatric conditions.

### Limitations of study

We would like to highlight several limitations of the current study that may provide avenues for future investigations. One is our reliance on c-fos as a neural activity marker for our regional ensemble analyses, as other IEGs may better capture activation in certain cell types throughout the brain^85^. Another limitation is that our sequencing study included only proestrus females; thus, we were unable to detect any sex- or estrous cycle-driven transcriptional signatures that may prime LS*^Nts-Sst^*neurons for ensemble recruitment. Lastly, despite our best efforts to employ COnFOn viruses to manipulate LS*^Nts-Sst^* neurons, we were unable to attain sufficient viral expression to establish the causal link between LS*^Nts-Sst^* neuron activity and our behavioral effects. Despite these limitations, we believe this work provides a rigorous and convincing description of a novel mechanism whereby cycling reproductive hormones can alter fundamental memory processes.

## AUTHOR CONTRIBUTIONS

NEB: Conceptualization, Data Curation, Formal Analysis, Investigation, Methodology, Software, Visualization, Writing – Original Draft, Writing – Reviewing and Editing; PJT: Data Curation, Formal Analysis, Investigation, Methodology, Software, Visualization, Writing – Reviewing and Editing; NSD: Formal Analysis, Investigation, Writing – Reviewing and Editing; KHAB: Formal Analysis, Investigation, Writing – Reviewing and Editing; GCB: Formal Analysis, Investigation, Writing – Reviewing and Editing; ASD: Formal Analysis, Investigation, Writing – Reviewing and Editing; GCN: Formal Analysis, Investigation, Writing – Reviewing and Editing; MP: Formal Analysis, Investigation, Writing – Reviewing and Editing; TRH: Data Curation, Formal Analysis, Investigation, Methodology, Software, Visualization, Writing – Reviewing and Editing; EKL: Conceptualization, Data Curation, Formal Analysis, Funding Acquisition, Investigation, Methodology, Project Administration, Resources, Supervision, Visualization, Writing – Reviewing and Editing

## ACKNOWLEDGEMENTS

Many thanks are due to Dr. Jeremy Simon for initial bioinformatics processing, Drs. Andrew Hardaway and Meghan Flannigan for guidance with fiber photometry analysis pipelines, Drs. Zoe McElligot and Gina Leinninger for sharing the *Nts*-Cre mouse line, and to Dara Russell, Elazar Barrett, Mandy Biraud, Amy Farthing, Alexa Jo Tellez, Bryana Whitaker Hardin, and May Rudd for assistance with animal husbandry and estrous cycle monitoring.

## EXPERIMENTAL MODEL AND STUDY PARTICIPANT DETAILS

Adult (4-8 months of age) male and female C57Bl/6J (#000664, Jackson Laboratories, Bar Harbor, ME, USA) and double heterozygous SST-IRES-FlpO (#031629)^93^ x NTS-Cre (#017525)^90^ mice (with a C57Bl/6J background) were bred in house for the purposes of this study. Prior to behavioral testing, mice were housed 2-6 mice per cage with *ad libitum* access to food and water on a 12-hour light-dark cycle. For experiments involving threat memory recall, mice were single-housed starting the first day of handling and remained single-housed for the duration of the behavioral experiment. All experiments were approved in advance by the Institutional Animal Care and Use Committees at North Carolina State University and University of Alabama at Birmingham and were conducted in accordance with the National Institutes of Health’s Guide for the Care and Use of Laboratory Animals.

## METHOD DETAILS

### Estrous cycle monitoring

Mice received daily vaginal (female) or sham (male) lavages between 8:00-9:00am starting 8 days prior to behavioral testing and continued throughout the duration of the experiment, as previously described^29^. Cells were stained using hematoxylin and eosin and characterized according to standard practices^94^. Proestrus was determined by a predominance of nucleated epithelial cells, estrus by a predominance of cornified epithelial cells, and diestrus by the presence of leukocytes. Mice that exhibited irregular estrous cycling (defined by ≥ 5 consecutive days in the same stage) were excluded from analyses.

### Viral Vectors

Viral vectors were purchased from Addgene (Watertown, MA, USA) and included AAV9-CaMKIIa-hM3D(Gq)-mCherry (#50476-AAV9)^86^, AAV9-CaMKIIa-hM4D(Gi)-mCherry (#50477-AAV9)^87^, AAV9-CaMKII-GCaMP6f (#100834-AAV9)^88^, AAV8-Ef1a-Con/Fon-mCherry (#137132-AAV8)^61^, and AAV8-Ef1a-Con/Fon-GCaMP6f (#137122-AAV8)^61^.

### Stereotaxic surgeries

Mice were anesthetized using isoflurane (5% for induction, 1-3% for maintenance) and placed in a stereotaxic frame (Stoelting, Wood Dale, IL, USA). Fur was removed, and the scalp was cleaned with Betadine and ethanol prior to a midline incision. The skull was cleaned using ethanol and hydrogen peroxide, and burr holes were made using a dental drill. Viral vectors were infused into the LS (AP +0.60, ML ±0.42, DV −3.20) at a rate of 1 nl/s using World Precision Instruments (Sarasota, FL, USA) digital injectors.

For pan-neuronal chemogenetic experiments, 0.15 μl of either AAV9-CaMKIIa-hM3D(G_q_)-mCherry or AAV9-CaMKIIa-hM4D(G_i_)-mCherry was bilaterally infused into the LS of C57Bl/6J mice. For Cre- and Flp-dependent fiber photometry experiments, 0.50 μl of AAV8-Ef1a-COnFOn-GCaMP6f was unilaterally infused into the right LS of double heterozygous *Nts*-Cre x *Sst*-FlpO mice. For Cre- and Flp-dependent anterograde tracing, 0.30 μl of AAV8-Ef1a-COnFOn-mCherry was bilaterally infused into the LS of double heterozygous *Nts*-Cre x *Sst*-FlpO mice, and brains were harvested 6 weeks later. For co-localization of LS*^Nts-Sst^* neurons with c-fos immunolabeling, 0.30 μl of AAV8-Ef1a-COnFOn-mCherry was bilaterally infused into the LS of female double heterozygous *Nts*-Cre x *Sst*-FlpO mice three weeks prior to behavioral testing.

For fiber photometry experiments, a fiber optic ferrule (200 μm core, 0.37 NA, 3.5 mm length; Newdoon, Hangzhou City, China) was implanted immediately following viral infusions at the same coordinates. Fibers were affixed to the skull with two anchor screws using C&B Metabond (Parkell, Brentwood, NY, USA) and dental cement (A-M Systems, Sequim, WA, USA). For all other surgeries, incision sites were closed using a surgical staple which was removed after 10 days. Mice received analgesia (Carprofen, 5 mg/kg) immediately prior to surgery and once daily for two days following surgery.

### Behavioral testing

<u>Handling.</u> Prior to behavioral testing, mice were handled by the experimenter for 3 minutes for at least 3 consecutive days in room adjacent to the behavioral testing area. For chemogenetic experiments, mice underwent 5 consecutive days of handling, with habituation to intraperitoneal injections using sterile saline immediately following handling on the second and fourth days. For fiber photometry experiments, mice were habituated to tethering and placed in a clean, lidless home cage while tethered to a patch cord for 10 minutes immediately following handling.

<u>Threat conditioning and recall.</u> Mice were randomly assigned to experimental groups and acclimated in an adjacent room for at least 30 minutes. Threat conditioning and recall were conducted in either Coulbourn Habitest (Harvard Bioscience Inc., Holliston, MA, USA) or Lafayette Instruments (Lafayette, IN, USA) modular operant chambers housed within sound-attenuating cubicles (Lafayette Instruments). Cued threat conditioning consisted of a 240 s baseline period followed by six co-terminating pairings of the CS (20 s, 5 kHZ, 75 dB pure tone) with the US (2 s, 0.5 mA foot shock) with 100 s ITIs. The conditioning context consisted of two clear plexiglass walls and two stainless steel walls with an aluminum shock grid floor and 70% EtOH was used as a contextual odorant. Cued recall consisted of a 240 s baseline period followed by ten presentations of the CS with 100 s ITIs. The recall context consisted of curved black plexiglass walls and black plexiglass flooring with isopropanol as a contextual odorant. CS-only groups experienced six presentations of the CS in the conditioning context and underwent cued recall identical to the threat conditioned groups. Context-forward threat conditioning was performed in the same context as cued threat conditioning and consisted of a 240 s baseline period followed by six presentations of the US with 100 s ITIs in the same context. Contextual recall consisted of 180 s in the conditioning context. Mice assigned to naïve control groups were placed in a clean cage and sacrificed at the same timepoint as conditioned mice for a given experiment. Chambers were cleaned with chlorohexidine between animals. Freezing was quantified using FreezeFrame V5 (Actimetrics, Wilmette, IL, USA). Freezing thresholds were determined for each animal based on the highest motion index for which the mouse displayed no movement except necessary for breathing for ≥ 3 s, unless otherwise stated. The percent of time spent freezing was analyzed between groups across trials during conditioning using a two-way repeated measures ANOVA. The percent of time spent freezing during the baseline period and the average time spent freezing across all CS presentations during recall testing were analyzed between groups using a one-way ANOVA. If group differences were observed for freezing during the baseline period of recall testing, those values were subtracted from the average CS freezing for each animal to account for contextual generalization between groups, as previously described^95^. Although freezing is the predominant behavioral response rodents display to threats^35,36^, including C57Bl/6J mice of both sexes^29^, others have demonstrated sex differences in alternative conditioned responses to threats^96–98^, prompting us to also quantify shock reactivity and active conditioned responses. Shock reactivity was determined by the FreezeFrame motion index^99^. Active behavioral responses during conditioning and recall were manually scored and expressed as flight scores, defined as the number of darts, jumps, and tail rattles per minute, as previously described^29^.

<u>Elevated plus maze.</u> Mice were acclimated in an adjacent room for at least 30 minutes prior to testing. The plus maze consisted of two open arms (30 cm) and two closed arms (30 cm with 15 cm walls) raised 40 cm from the table surface (Panlab, Barcelona, Spain). Each mouse was placed in the center of the plus maze and allowed to explore for 5 minutes. Behavior was recorded with an overhead camera and analyzed using AnyMaze software (Stoelting). Percent open-arm entries was calculated by dividing the number of open arm entries by the total number of entries into the open and closed arms and multiplying by 100.

### Immunohistochemistry and c-fos quantification following conditioning and recall

Mice were anesthetized with tribromoethanol (TBE) and transcardially perfused with phosphate buffered saline (PBS) and 4% paraformaldehyde (PFA) 90 minutes after entering the operant chamber or clean cage (naïve treatment). Brains were postfixed overnight in 4% PFA, washed with PBS, and cryoprotected in 30% sucrose in PBS. Brains were then embedded in a 2:1 mixture of OCT Compound (ThermoFisher Scientific, Waltham, MA, USA) and 30% sucrose in PBS and stored at −80°C until cryosectioning. Coronal sections were collected at at 50 μm thickness in 300 μm intervals spanning from approximately +2.95 to −5.80 AP relative to bregma for immunofluorescence staining as previously described^42^. Antibodies used included rabbit anti-c-fos primary antibody (Synaptic Systems # 226008, Goettingen, Germany; 0.25 μg/mL dilution) and donkey anti-rabbit 647 secondary antibody (Jackson ImmunoResearch #711605152; West Grove, PA, USA; 1.25 μg/mL dilution). Following immunostaining, sections were counterstained with DAPI (Invitrogen, Carlsbad, CA, USA), mounted onto slides and coverslipped with Prolong Gold Antifade Mountant (ThermoFisher Scientific). Slides were stored at 4°C until imaging. For post-conditioning analyses, all sections were imaged on an Olympus FV3000 confocal microscope (Olympus Global, Tokyo, Japan) at 20x magnification on a single z-plane focused at 10 μm tissue depth with uniform settings obtained from optimization on a representative conditioned proestrus brain. For post-recall analyses, sections were imaged on an Olympus VS200 Slide Scanner microscope at 20x magnification focused at maximum DAPI intensity. The ImageJ plug-in ABBA^85^ was used to register regions defined by the Allen Brain Atlas to each experimental brain, and c-fos positive cells were quantified from brain regions previously observed to be engaged in both sexes following threat conditioning^42^. Images were despeckled and thresholded to twice the background fluorescence intensity prior to manual quantification of c-fos positive cells in ImageJ. Cell counts were normalized to area (mm^2^), and values for regions spanning multiple brain sections and/or hemispheres were averaged together to obtain a single value per region for each brain.

### Network analyses

Functional connectivity and network analyses were conducted for each experimental group as previously described^42^. Correlation matrices were constructed using Pearson’s correlation coefficient for all pairwise comparisons of regional c-fos counts for all 114 brain regions following threat conditioning or naïve treatment. Networks of correlated interregional c-fos counts were constructed within each threat conditioned group using a *p* value cut off of 0.02, corresponding to a Pearson’s Correlation coefficient *r* ≥ 0.98. Only positive correlations were included. Clusters of nodes within each network were created using Markov clustering^100^, and degree and betweenness were calculated for each node. Hub regions were identified as nodes in the top 20 percentile for both degree (number of edges) and betweenness (number of shortest paths through a node). Sequential *p*-value pruning was used to identify hub regions most reliably appearing as network hubs across 10 different *p*-value thresholds (0.005 to 0.05). All analyses were conducted in Python using the Brain Connectivity Toolbox^101^. Code used for network analyses and visualization is available on Github.

### Anterograde tracing analyses

Brains were harvested and cryosectioned as described above at 50 μm thickness. Sections were collected at 300 μm intervals spanning from approximately +3.20 to −6.70 AP relative to Bregma and counter-stained with DAPI for brain-wide survey of mCherry terminals. For any brain region displaying terminals in at least four different brains, 20x magnification images were obtained on an Olympus FV3000 confocal microscope with uniform settings determined using a representative female brain. Serial z-stacked images spanning the entire thickness of the tissue were flattened into a maximum intensity projection for quantification of terminal densities. For each region of interest, background threshold was manually determined, and the number of pixels above the threshold was quantified using the Analyze Particles function in ImageJ. Pixel counts were normalized to area (mm^2^), then divided by the number of total starter cells (number of mCherry somas in the LS) for each brain to determine the normalized mCherry density for each region of interest.

### Chemogenetics

C57Bl/6J mice received bilateral infusions of either an inhibitory (hM4D) or excitatory (hM3D) DREADD directed to the LSr. Three weeks later, mice underwent auditory threat conditioning and cued recall following intraperitoneal injections of either the DREADD activator DCZ (0.1 mg/kg dissolved in 0.1% DMSO in saline^89^; MedChemExpress, Monmouth Junction, NJ, USA) or a vehicle injection consisting of 0.1% DMSO in saline. For hM4D experiments, intraperitoneal injections occurred 30 minutes prior to entering the operant chambers, and mice were euthanized via transcardial perfusion 90 minutes after entering the operant chamber for cued recall. For hM3D experiments, injections occurred one hour prior to entering the operant chambers. Several days following cued recall, mice used in hM3D experiments received an additional intraperitoneal injection of either DCZ or vehicle and were euthanized via transcardial perfusion two hours later. A subset of brains from each experimental group was sectioned and processed for immunostaining for c-fos as described above to validate DREADD efficacy. Sections were imaged on an Olympus FV3000 confocal microscope at 20x magnification on a single z-plane focused at 10 μm tissue depth with uniform settings obtained from optimization on a representative DCZ-treated brain. Images were analyzed in QuPath software^91^ to perform cell detection in the DAPI channel and positive cell detection of c-fos and mCherry were considered when cells expressed twice the background maximum fluorescence in that channel.

### Single nucleus RNA sequencing

<u>Tissue collection, nuclei isolation, and RNA sequencing.</u> C57Bl/6J female mice in proestrus were randomly assigned to threat conditioned or naïve control groups (n=4/group). Five minutes after exiting the conditioning chamber, mice were anesthetized with TBE and euthanized via rapid decapitation. Brains were removed within two minutes, submerged in ice-cold Hibernate® AB Complete (Transnetyx, Cordova, TN, USA) for 30 s, and sectioned at 1 mm using a chilled stainless-steel brain matrix (Zivic Instruments, Fremont, CA, USA). LS punches (1.25 mm, Stoelting, Wood Dale, IL, USA) were collected under a dissecting microscope, snap frozen on dry ice, and stored at −80 °C. LS punches were combined into one sample for each experimental group prior to nuclei isolation following Phillips *et al*. (2022)^102^ with modifications. Samples were thawed on wet ice for 5 min prior to the addition of lysis buffer (10 mM Tris-HCl pH 7.4, 10 mM NaCl, 3 mM MgCl₂, 0.1% Nonidet P40 in nuclease-free ddH₂O). After 15 min on ice with periodic swirling, 1 mL Hibernate® AB was added to quench the reaction, and the suspension was triturated through decreasing fire-polished pipette diameters. The lysate was filtered (35 µm), centrifuged (500 G, 10 min, 4 °C) and resuspended twice for washing, then resuspended in 500 µL Hibernate® AB. Nuclei quality and quantity was verified with trypan blue staining. Nuclei were labeled with 7-AAD (Thermo Scientific; 5 µL/million nuclei), and singlets were sorted with a SH800Z Cell Sorter (Sony Biotechnology, San Jose, CA, USA) into BSA-coated tubes. The Advanced Analytics Core at the University of North Carolina Chapel Hill (UNC) processed the suspensions (1000 nuclei/µL) for the 10x Chromium Next GEM Single Cell Multiome ATAC + Gene Expression Kit (10x Genomics, Pleasanton, CA, USA). For each sample, the maximum number of nuclei were loaded in each inlet with two inlets per group to target recovery of 20,000 nuclei per group. The UNC High-Throughput Sequencing Facility sequenced gene expression libraries on an Illumina NovaSeq 6000 Sequencer using v1.5 chemistry (28 × 10 × 10 × 90 cycles) to target >50,000 reads/nucleus.

<u>Preprocessing and quality control.</u> Reads were processed with CellRanger-ARC v7.0.0 (10x Genomics) against the mouse reference transcriptome (refdata-gex-mm10-2020-A). Each replicate produced filtered and unfiltered AnnData H5 objects. All quality control and subsequent analyses were performed using R v4.4.1 (The R Foundation, Vienna, Austria) and Python v3.10.16 (The Python Software Foundation, Beaverton, OR, USA) within Conda v24.11.3 (Anaconda Inc., Austin, TX, USA) via Miniforge (conda-forge, https://conda-forge.org). Background RNA contamination was estimated and corrected with SoupX v1.6.2^103^, and low-quality nuclei were removed if mitochondrial transcripts (prefix “^mt-”) exceeded 5%. Likely doublets were identified with scDblFinder v1.20.2^104^. Initial clustering of all replicates used Seurat v5.3.0^105^. Cluster markers and reference-based annotations were assigned via MapMyCells (RRID:SCR_024672) using the Allen Institute/10x Genomics CCN20230722 “Whole Mouse Brain” taxonomy (Allen Institute for Brain Science, 2004, 2011). Clusters absent from the LS in the ABC Atlas^106^ or lacking biologically relevant markers or MapMyCells class identity were excluded. The final post-processing nuclei count was 13,457 and 11,455 for the conditioned and naïve groups, respectively.

<u>Latent Representation and Probabilistic Analysis.</u> Preliminary Seurat clustering revealed strong batch-dependent effects. To correct these, we trained a variational autoencoder with scVI (unsupervised step) and scANVI (fine-tuning step) (scvi-tools v1.4.0^58^) using a modified implementation to expose and automatically record the evidence lower bound (ELBO), reconstruction loss, and Kullback–Leibler (KL) divergence at the end of each training and fine-tuning epoch. The scVI model incorporated 2 hidden layers (128 units), 34-dimensional latent space, batch size 500, dropout 0.2, learning rate 5 × 10−4, weight decay 1 × 10−4, 175 epochs, and KL-warmup 50 epochs. Batch and condition covariates were one-hot encoded within the decoder to prevent embedding of technical effects. scANVI fine-tuning used MapMyCells supertypes (learning rate 5 × 10−5, weight decay 1 × 10−5, 75 epochs). Leiden clustering (cosine metric, k = 45, resolution 0.8) and UMAP embedding (spread 1.1, min.dist 0.2) were performed. Biologically identical clusters were merged manually before differential expression was calculated with scvi-tools (mode = “change,” δ = 0.25, 5,000 Monte Carlo samples). Comparisons included each cluster versus all others, each cluster between conditions, and pairwise cluster-wise analyses. For all, FDR ≤ 0.05 was considered significant. Data visualization used ggplot2 v3.5.2 in R and anndata (v1.16.0), scanpy (v1.11.1), pandas, numpy (v2.1.3), scipy, igraph, leidenalg, and h5py in Python.

### Multiplex fluorescent *in situ* hybridization

C57Bl/6J male and female mice targeted to either proestrus or diestrus were randomly assigned to threat conditioned or naïve control groups. Five minutes after exiting the conditioning chamber, mice were anesthetized with TBE and euthanized via rapid decapitation. Brains were flash frozen in 2-methylbutane and stored at −80°C. Brains were then cryosectioned at 15 μm thickness directly onto SuperFrost Plus microscope slides (Fisher Scientific), and slides were stored at −80°C in sealed bags. Sections containing the LS were processed for fluorescent *in situ* hybridization using the RNAScope Multiplex Fluorescent kit (ACD Bio; Newark, California, USA) following manufacturer instructions with probes targeting *Sst* (404631-C1), *Nts* (420441-C2), *Crhr2* (413201-C3), and *Fos* (316921-C4). Probes were fluorescently tagged for imaging using Opal 520 (FP1487001KT; Akoya Biosciences; Marlborough, MA, USA), Opal 570 (FP1488001KT), Opal 650 (FP1496001KT), and Opal 690 (FP1497001KT) reagents and counterstained with DAPI.

Sections were imaged within 48 hours on an Olympus FV3000 confocal microscope at 20x magnification using standardized settings obtained from optimization on a representative conditioned proestrus sample. Images were analyzed in QuPath software^91^ to perform cell detection in the DAPI channel and cell type classification in each probe channel using thresholds obtained from negative control probe staining for each respective brain.

### Fiber photometry

Three weeks following stereotaxic infusions of a calcium sensor and implantation of a fiber optic ferrule in the LS, calcium fluorescent activity was measured throughout cued threat conditioning and recall testing using a Neurophotometrics (San Diego, CA, USA) FP3002 system connected via Doric (Quebec, Canada) fiber-optic patch cords. Fluorescence in the calcium indicator channel (450 nm) and the isosbestic channel (415 nm) were collected at a rate of 40 Hz in an alternating pattern, with laser intensity set to 50 mW for each respective channel.

Analysis of fiber photometry signals was performed in MATLABR2022 (MathWorks Inc, Natick, MA, USA) using custom code modified from Neurophotometrics suggested code (available on www.neurophotometrics.com/documentation). Signals from the calcium indicator channel and the isosbestic channel were deinterleaved. To correct for photobleaching, the isosbestic signal was fitted with a bioexponential curve and linearly scaled to fit the calcium indicator signals. Calcium indicator signals were normalized to the scaled fitted curves, and z-scores were calculated from the normalized data to quantify calcium changes in fluorescence over time. To quantify fluorescent changes surrounding CS presentations, z-scores were normalized to a 5-second baseline period preceding the event. AUC analyses were performed using MATLAB trapz function. To analyze the change in AUC across conditioning, we calculated the slope of AUC for specified periods across CS-US pairings. For within-subject comparisons between AUC slopes during conditioning and cued freezing during recall, we performed simple linear regressions for each experimental group. For within subject comparisons of calcium activity and freezing during cued recall, we calculated the Pearson’s R coefficient for each animal and compared R coefficients between groups using a one-way ANOVA.

### Estradiol administration

Three weeks following stereotaxic infusions of a viral reporter, female mice in diestrus underwent auditory threat conditioning as described above. Four hours prior to conditioning, mice received a subcutaneous injection of either 17β-Estradiol (Sigma Aldrich; 0.2mg/kg in peanut oil) or vehicle (peanut oil). Ninety-minutes after conditioning, mice were anesthetized with TBE and perfused. Brains were sectioned and processed for immunostaining for c-fos as described above for chemogenetics. Sections were imaged on an Olympus FV3000 confocal microscope at 20x magnification on a single z-plane focused at 10 μm tissue depth with uniform settings obtained from optimization on a representative estradiol-treated brain. Cell counts for mCherry+, c-fos+, and double-labelled cells were manually collected using ImageJ as described above.

### Rigor and reproducibility

All behavioral tests were conducted between 2:00-6:00pm to target circadian surges of hormone levels in proestrus prior to the dark phase^107^. Female mice were tested on the day of proestrus or the first day of diestrus. Male mice were yoked to female mice of different experimental groups, and testing schedules were matched for yoked pairs. Experimental groups were randomly assigned and balanced across cohorts for all experiments. Researchers were blind to experimental group during data analyses as much as possible. Surgical targeting was verified for each experiment, and animals were excluded from all analyses if substantial viral expression or ferrule placement was detected outside of the target region.

## QUANTIFICATION AND STATISTICAL ANALYSIS

Unless otherwise specified in legends, graphs represent group means ± standard error of the mean. Network analyses were performed using Python as described above. snSEQ data were analyzed using Python and R as described above. Fiber photometry data were analyzed using MATLAB as described above. All remaining statistics were performed using IBM SPSS Statistics software and graphed using GraphPad Prism 10. Normality of datasets were first confirmed using the Shapiro–Wilk test. Comparisons of group means were conducted using one- or two-way ANOVAs with and without repeating measures as appropriate. Greenhouse-Geisser corrections were applied for repeated-measures ANOVAs found to violate the assumption of sphericity via Mauchly’s test. Family-wise Bonferroni adjustments for post-hoc analyses were performed for three or more groups following a significant main effect or interactive effect from the parent ANOVA. Statistical tests are detailed in the Supplemental Tables.

## REFERENCES

1 Breslau, N., Davis, G. C., Andreski, P., Peterson, E. L. & Schultz, L. R. Sex differences in posttraumatic stress disorder. Arch Gen Psychiatry 54, 1044–1048 (1997). 10.1001/archpsyc.1997.01830230082012

2 Kilpatrick, D. G. et al. National estimates of exposure to traumatic events and PTSD prevalence using DSM-IV and DSM-5 criteria. J Trauma Stress 26, 537–547 (2013). 10.1002/jts.21848

3 Nillni, Y. I., Rasmusson, A. M., Paul, E. L. & Pineles, S. L. The Impact of the Menstrual Cycle and Underlying Hormones in Anxiety and PTSD: What Do We Know and Where Do We Go From Here? Curr Psychiatry Rep 23, 8 (2021). 10.1007/s11920-020-01221-9

4 Bryant, R. A. et al. The association between menstrual cycle and traumatic memories. J Affect Disord 131, 398–401 (2011). 10.1016/j.jad.2010.10.049

5 Soni, M., Curran, V. H. & Kamboj, S. K. Identification of a narrow post-ovulatory window of vulnerability to distressing involuntary memories in healthy women. Neurobiol Learn Mem 104, 32–38 (2013). 10.1016/j.nlm.2013.04.003

6 Franke, L. K. et al. Estradiol during (analogue-)trauma: Risk- or protective factor for intrusive re-experiencing? Psychoneuroendocrinology 143, 105819 (2022). 10.1016/j.psyneuen.2022.105819

7 Rieder, J. K., Kleshchova, O. & Weierich, M. R. Estradiol, stress reactivity, and daily affective experiences in trauma-exposed women. Psychol Trauma 14, 738–746 (2022). 10.1037/tra0001113

8 Nillni, Y. I. et al. Menstrual cycle effects on psychological symptoms in women with PTSD. J Trauma Stress 28, 1–7 (2015). 10.1002/jts.21984

9 Glover, E. M. et al. Inhibition of fear is differentially associated with cycling estrogen levels in women. J Psychiatry Neurosci 38, 341–348 (2013). 10.1503/jpn.120129

10 Mueller, J. M. et al. Dynamic community detection reveals transient reorganization of functional brain networks across a female menstrual cycle. Netw Neurosci 5, 125–144 (2021). 10.1162/netn_a_00169

11 Pritschet, L. et al. Functional reorganization of brain networks across the human menstrual cycle. Neuroimage 220, 117091 (2020). 10.1016/j.neuroimage.2020.117091

12 Woolley, C. S., Gould, E., Frankfurt, M. & McEwen, B. S. Naturally occurring fluctuation in dendritic spine density on adult hippocampal pyramidal neurons. J Neurosci 10, 4035–4039 (1990). 10.1523/JNEUROSCI.10-12-04035.1990

13 Woolley, C. S. & McEwen, B. S. Estradiol mediates fluctuation in hippocampal synapse density during the estrous cycle in the adult rat. J Neurosci 12, 2549–2554 (1992). 10.1523/JNEUROSCI.12-07-02549.1992

14 Warren, S. G., Humphreys, A. G., Juraska, J. M. & Greenough, W. T. LTP varies across the estrous cycle: enhanced synaptic plasticity in proestrus rats. Brain Res 703, 26–30 (1995). 10.1016/0006-8993(95)01059-9

15 Scharfman, H. E., Mercurio, T. C., Goodman, J. H., Wilson, M. A. & MacLusky, N. J. Hippocampal excitability increases during the estrous cycle in the rat: a potential role for brain-derived neurotrophic factor. J Neurosci 23, 11641–11652 (2003). 10.1523/JNEUROSCI.23-37-11641.2003

16 Jaric, I., Rocks, D., Greally, J. M., Suzuki, M. & Kundakovic, M. Chromatin organization in the female mouse brain fluctuates across the oestrous cycle. Nat Commun 10, 2851 (2019). 10.1038/s41467-019-10704-0

17 Rocks, D. et al. Sex-specific multi-level 3D genome dynamics in the mouse brain. Nat Commun 13, 3438 (2022). 10.1038/s41467-022-30961-w

18 Woolley, C. S. & McEwen, B. S. Roles of estradiol and progesterone in regulation of hippocampal dendritic spine density during the estrous cycle in the rat. J Comp Neurol 336, 293–306 (1993). 10.1002/cne.903360210

19 Freeman, M. E. in The Physiology of Reproduction, Vol. 2 (eds E. Knobil & J.D. Neill) Ch. 46, 613–658 (Raven Press, Ltd, 1994).

20 Viau, V. & Meaney, M. J. Variations in the hypothalamic-pituitary-adrenal response to stress during the estrous cycle in the rat. Endocrinology 129, 2503–2511 (1991). 10.1210/endo-129-5-2503

21 Eckel, L. A. & Geary, N. Endogenous cholecystokinin’s satiating action increases during estrus in female rats. Peptides 20, 451–456 (1999). 10.1016/s0196-9781(99)00025-x

22 Clegg, D. J. et al. Estradiol-dependent decrease in the orexigenic potency of ghrelin in female rats. Diabetes 56, 1051–1058 (2007). 10.2337/db06-0015

23 Applebey, S. V. et al. Characterizing Brainstem GLP-1 Control of Sensory-Specific Satiety in Male and Female Rats Across the Estrous Cycle. Biol Psychiatry 98, 249–259 (2025). 10.1016/j.biopsych.2025.01.012

24 Blaustein, J. D. in Hormones, brain and behavior (ed A. P. Arnold D. W. Pfaff, A. M. Etgen, S. E. Fahrbach, & R. T. Rubin) 67–107 (Elsevier Academic Press., 2009).

25 Blaustein, J. D. & Wade, G. N. Ovarian influences on the meal patterns of female rats. Physiol Behav 17, 201–208 (1976). 10.1016/0031-9384(76)90064-0

26 Anantharaman-Barr, H. G. & Decombaz, J. The effect of wheel running and the estrous cycle on energy expenditure in female rats. Physiol Behav 46, 259–263 (1989). 10.1016/0031-9384(89)90265-5

27 Milad, M. R. & Quirk, G. J. Fear extinction as a model for translational neuroscience: ten years of progress. Annu Rev Psychol 63, 129–151 (2012). 10.1146/annurev.psych.121208.131631

28 Bauer, E. P. Sex differences in fear responses: Neural circuits. Neuropharmacology 222, 109298 (2023). 10.1016/j.neuropharm.2022.109298

29 Ryherd, G. L., Bunce, A. L., Edwards, H. A., Baumgartner, N. E. & Lucas, E. K. Sex differences in avoidance behavior and cued threat memory dynamics in mice: Interactions between estrous cycle and genetic background. Horm Behav 156, 105439 (2023). 10.1016/j.yhbeh.2023.105439

30 Le Roux, C. E., Farthing, A. L. & Lucas, E. K. Dietary phytoestrogens recalibrate socioemotional behavior in C57Bl/6J mice in a sex- and timing-dependent manner. Horm Behav 168, 105678 (2025). 10.1016/j.yhbeh.2025.105678

31 Girden, E. & Culler, E. Conditioned responses in curarized striate muscle in dogs. Journal of Comparative Psychology 23, 261–274 (1937). 10.1037/h0058634

32 Radulovic, J., Jovasevic, V. & Meyer, M. A. Neurobiological mechanisms of state-dependent learning. Curr Opin Neurobiol 45, 92–98 (2017). 10.1016/j.conb.2017.05.013

33 Jovasevic, V. et al. GABAergic mechanisms regulated by miR-33 encode state-dependent fear. Nat Neurosci 18, 1265–1271 (2015). 10.1038/nn.4084

34 Meyer, M. A. A. et al. Neurobiological correlates of state-dependent context fear. Learn Mem 24, 385–391 (2017). 10.1101/lm.045542.117

35 Blanchard, R. J. & Blanchard, D. C. Crouching as an index of fear. J Comp Physiol Psychol 67, 370–375 (1969). 10.1037/h0026779

36 Fanselow, M. S. Conditioned and unconditional components of post-shock freezing. Pavlov J Biol Sci 15, 177–182 (1980). 10.1007/BF03001163

37 Botta, P. et al. Regulating anxiety with extrasynaptic inhibition. Nat Neurosci 18, 1493–1500 (2015). 10.1038/nn.4102

38 Blair, R. S. et al. Estrous cycle contributes to state-dependent contextual fear in female rats. Psychoneuroendocrinology 141, 105776 (2022). 10.1016/j.psyneuen.2022.105776

39 Josselyn, S. A., Kohler, S. & Frankland, P. W. Finding the engram. Nat Rev Neurosci 16, 521–534 (2015). 10.1038/nrn4000

40 Willett, J. A. et al. The estrous cycle modulates rat caudate-putamen medium spiny neuron physiology. Eur J Neurosci 52, 2737–2755 (2020). 10.1111/ejn.14506

41 Blume, S. R. et al. Sex- and Estrus-Dependent Differences in Rat Basolateral Amygdala. J Neurosci 37, 10567–10586 (2017). 10.1523/JNEUROSCI.0758-17.2017

42 du Plessis, K. C., Basu, S., Rumbell, T. H. & Lucas, E. K. Sex-Specific Neural Networks of Cued Threat Conditioning: A Pilot Study. Front Syst Neurosci 16, 832484 (2022). 10.3389/fnsys.2022.832484

43 Spiegel, E. A., Miller, H. R. & Oppenheimer, M. J. Forebrain and rage reactions. Journal of Neurophysiology 3, 538–548 (1940).

44 Slotnick, B. M., McMullen, M. F. & Fleischer, S. Changes in emotionality following destruction of the septal area in albino mice. Brain Behav Evol 8, 241–252 (1973). 10.1159/000124357

45 Besnard, A. et al. Dorsolateral septum somatostatin interneurons gate mobility to calibrate context-specific behavioral fear responses. Nat Neurosci 22, 436–446 (2019). 10.1038/s41593-018-0330-y

46 Besnard, A., Miller, S. M. & Sahay, A. Distinct Dorsal and Ventral Hippocampal CA3 Outputs Govern Contextual Fear Discrimination. Cell Rep 30, 2360–2373 e2365 (2020). 10.1016/j.celrep.2020.01.055

47 Opalka, A. N. & Wang, D. V. Hippocampal efferents to retrosplenial cortex and lateral septum are required for memory acquisition. Learn Mem 27, 310–318 (2020). 10.1101/lm.051797.120

48 Hashimoto, M. et al. Lateral septum modulates cortical state to tune responsivity to threat stimuli. Cell Rep 41, 111521 (2022). 10.1016/j.celrep.2022.111521

49 Chen, M., Li, J., Shan, W., Yang, J. & Zuo, Z. Auditory fear memory retrieval requires BLA-LS and LS-VMH circuitries via GABAergic and dopaminergic neurons. EMBO Rep 26, 1816–1834 (2025). 10.1038/s44319-025-00403-x

50 Calandreau, L., Jaffard, R. & Desmedt, A. Dissociated roles for the lateral and medial septum in elemental and contextual fear conditioning. Learn Mem 14, 422–429 (2007). 10.1101/lm.531407

51 Sheehan, T. P., Chambers, R. A. & Russell, D. S. Regulation of affect by the lateral septum: implications for neuropsychiatry. Brain Res Brain Res Rev 46, 71–117 (2004). 10.1016/j.brainresrev.2004.04.009

52 Jacobs, N. S., Cushman, J. D. & Fanselow, M. S. The accurate measurement of fear memory in Pavlovian conditioning: Resolving the baseline issue. J Neurosci Methods 190, 235–239 (2010). 10.1016/j.jneumeth.2010.04.029

53 Turrero Garcia, M., et al. Transcriptional profiling of sequentially generated septal neuron fates. Elife 10 (2021). 10.7554/eLife.71545

54 Phillips, R. A., 3rd et al. Transcriptomic characterization of human lateral septum neurons reveals conserved and divergent marker genes across species. iScience 28, 111820 (2025). 10.1016/j.isci.2025.111820

55 Wang, Y. et al. Spatial Cell Atlas of Lateral Septum Reveals Changes Underlying Anxiety and Fear Learning Deficits in Mice with Abnormal Immunity. Int J Biol Sci 21, 6389–6410 (2025). 10.7150/ijbs.117249

56 Chen, G. et al. Cellular and circuit architecture of the lateral septum for reward processing. Neuron 112, 2783–2798 e2789 (2024). 10.1016/j.neuron.2024.06.004

57 Simon, R. C. et al. Opioid-driven disruption of the septum reveals a role for neurotensin-expressing neurons in withdrawal. Neuron 113, 2325–2343 e2329 (2025). 10.1016/j.neuron.2025.04.024

58 Lopez, R., Regier, J., Cole, M. B., Jordan, M. I. & Yosef, N. Deep generative modeling for single-cell transcriptomics. Nat Methods 15, 1053–1058 (2018). 10.1038/s41592-018-0229-2

59 Azevedo, E. P. et al. A limbic circuit selectively links active escape to food suppression. Elife 9 (2020). 10.7554/eLife.58894

60 Anthony, T. E. et al. Control of stress-induced persistent anxiety by an extra-amygdala septohypothalamic circuit. Cell 156, 522–536 (2014). 10.1016/j.cell.2013.12.040

61 Fenno, L. E. et al. Comprehensive Dual- and Triple-Feature Intersectional Single-Vector Delivery of Diverse Functional Payloads to Cells of Behaving Mammals. Neuron 107, 836–853 e811 (2020). 10.1016/j.neuron.2020.06.003

62 Chen, Z. et al. A circuit from lateral septum neurotensin neurons to tuberal nucleus controls hedonic feeding. Mol Psychiatry 27, 4843–4860 (2022). 10.1038/s41380-022-01742-0

63 Li, L. et al. Social trauma engages lateral septum circuitry to occlude social reward. Nature 613, 696–703 (2023). 10.1038/s41586-022-05484-5

64 An, M. et al. Lateral Septum Somatostatin Neurons are Activated by Diverse Stressors. Exp Neurobiol 31, 376–389 (2022). 10.5607/en22024

65 McCarthy, M. M., Arnold, A. P., Ball, G. F., Blaustein, J. D. & De Vries, G. J. Sex differences in the brain: the not so inconvenient truth. J Neurosci 32, 2241–2247 (2012). 10.1523/JNEUROSCI.5372-11.2012

66 Contreras, C. M. & Gutierrez-Garcia, A. G. Estrogen and progesterone priming induces lordosis in female rats by reversing the inhibitory influence of the infralimbic cortex on neuronal activity of the lateral septal nucleus. Neurosci Lett 732, 135079 (2020). 10.1016/j.neulet.2020.135079

67 Contreras, C. M., Molina, M., Saavedra, M. & Martinez-Mota, L. Lateral septal neuronal firing rate increases during proestrus-estrus in the rat. Physiol Behav 68, 279–284 (2000). 10.1016/s0031-9384(99)00169-9

68 Yiu, A. P. et al. Neurons are recruited to a memory trace based on relative neuronal excitability immediately before training. Neuron 83, 722–735 (2014). 10.1016/j.neuron.2014.07.017

69 Barth, C., Villringer, A. & Sacher, J. Sex hormones affect neurotransmitters and shape the adult female brain during hormonal transition periods. Front Neurosci 9, 37 (2015). 10.3389/fnins.2015.00037

70 Kim, H. J. J. et al. Impact of the mouse estrus cycle on cannabinoid receptor agonist-induced molecular and behavioral outcomes. Pharmacol Res Perspect 10, e00950 (2022). 10.1002/prp2.950

71 Spencer, J. L., Waters, E. M., Milner, T. A. & McEwen, B. S. Estrous cycle regulates activation of hippocampal Akt, LIM kinase, and neurotrophin receptors in C57BL/6 mice. Neuroscience 155, 1106–1119 (2008). 10.1016/j.neuroscience.2008.05.049

72 Woolley, C. S., Weiland, N. G., McEwen, B. S. & Schwartzkroin, P. A. Estradiol increases the sensitivity of hippocampal CA1 pyramidal cells to NMDA receptor-mediated synaptic input: correlation with dendritic spine density. J Neurosci 17, 1848–1859 (1997). 10.1523/JNEUROSCI.17-05-01848.1997

73 Foy, M. R. et al. 17beta-estradiol enhances NMDA receptor-mediated EPSPs and long-term potentiation. J Neurophysiol 81, 925–929 (1999). 10.1152/jn.1999.81.2.925

74 Shughrue, P. J., Lane, M. V. & Merchenthaler, I. Comparative distribution of estrogen receptor-alpha and -beta mRNA in the rat central nervous system. J Comp Neurol 388, 507–525 (1997). 10.1002/(sici)1096-9861(19971201)388:4<507::aid-cne1>3.0.co;2-6

75 Mitra, S. W. et al. Immunolocalization of estrogen receptor beta in the mouse brain: comparison with estrogen receptor alpha. Endocrinology 144, 2055–2067 (2003). 10.1210/en.2002-221069

76 Risold, P. Y. & Swanson, L. W. Chemoarchitecture of the rat lateral septal nucleus. Brain Res Brain Res Rev 24, 91–113 (1997). 10.1016/s0165-0173(97)00008-8

77 Ramirez, F., Moscarello, J. M., LeDoux, J. E. & Sears, R. M. Active avoidance requires a serial basal amygdala to nucleus accumbens shell circuit. J Neurosci 35, 3470–3477 (2015). 10.1523/JNEUROSCI.1331-14.2015

78 Zhou, J., Hormigo, S., Sajid, M. S. & Castro-Alamancos, M. A. Role of the Nucleus Accumbens in Signaled Avoidance Actions. eNeuro 11 (2024). 10.1523/ENEURO.0314-24.2024

79 Badarnee, M. et al. Intersect between brain mechanisms of conditioned threat, active avoidance, and reward. Commun Psychol 3, 32 (2025). 10.1038/s44271-025-00197-7

80 Nelson, R. J. & Kriegsfeld, L. J. in An introduction to behavioral endocrinology 275–333 (Sinauer Associates Inc. Publishers, 2017).

81 Pestana, J. E. & Graham, B. M. The impact of estrous cycle on anxiety-like behaviour during unlearned fear tests in female rats and mice: A systematic review and meta-analysis. Neurosci Biobehav Rev 164, 105789 (2024). 10.1016/j.neubiorev.2024.105789

82 Rocks, D., Cham, H. & Kundakovic, M. Why the estrous cycle matters for neuroscience. Biol Sex Differ 13, 62 (2022). 10.1186/s13293-022-00466-8

83 Feltenstein, M. W., Henderson, A. R. & See, R. E. Enhancement of cue-induced reinstatement of cocaine-seeking in rats by yohimbine: sex differences and the role of the estrous cycle. Psychopharmacology (Berl) 216, 53–62 (2011). 10.1007/s00213-011-2187-6

84 Johnson, A. R. et al. Cues play a critical role in estrous cycle-dependent enhancement of cocaine reinforcement. Neuropsychopharmacology 44, 1189–1197 (2019). 10.1038/s41386-019-0320-0

85 Chiaruttini, N. et al. ABBA+BraiAn, an integrated suite for whole-brain mapping, reveals brain-wide differences in immediate-early genes induction upon learning. Cell Rep 44, 115876 (2025). 10.1016/j.celrep.2025.115876

86 Chandrasekar, A. et al. Parvalbumin Interneurons Shape Neuronal Vulnerability in Blunt TBI. Cereb Cortex 29, 2701–2715 (2019). 10.1093/cercor/bhy139

87 Yang, T. et al. Social Control of Hypothalamus-Mediated Male Aggression. Neuron 95, 955–970 e954 (2017). 10.1016/j.neuron.2017.06.046

88 Kaufman, A. M., Geiller, T. & Losonczy, A. A Role for the Locus Coeruleus in Hippocampal CA1 Place Cell Reorganization during Spatial Reward Learning. Neuron 105, 1018–1026 e1014 (2020). 10.1016/j.neuron.2019.12.029

89 Nagai, Y. et al. Deschloroclozapine, a potent and selective chemogenetic actuator enables rapid neuronal and behavioral modulations in mice and monkeys. Nat Neurosci 23, 1157–1167 (2020). 10.1038/s41593-020-0661-3

90 Leinninger, G. M. et al. Leptin action via neurotensin neurons controls orexin, the mesolimbic dopamine system and energy balance. Cell Metab 14, 313–323 (2011). 10.1016/j.cmet.2011.06.016

91 Bankhead, P. et al. QuPath: Open source software for digital pathology image analysis. Sci Rep 7, 16878 (2017). 10.1038/s41598-017-17204-5

92 Traag, V. A., Waltman, L. & van Eck, N. J. From Louvain to Leiden: guaranteeing well-connected communities. Sci Rep 9, 5233 (2019). 10.1038/s41598-019-41695-z

93 He, M. et al. Strategies and Tools for Combinatorial Targeting of GABAergic Neurons in Mouse Cerebral Cortex. Neuron 91, 1228–1243 (2016). 10.1016/j.neuron.2016.08.021

94 Cora, M. C., Kooistra, L. & Travlos, G. Vaginal Cytology of the Laboratory Rat and Mouse: Review and Criteria for the Staging of the Estrous Cycle Using Stained Vaginal Smears. Toxicol Pathol 43, 776–793 (2015). 10.1177/0192623315570339

95 Baumgartner, N. E., Biraud, M. C. & Lucas, E. K. Sex differences in socioemotional behavior and changes in ventral hippocampal transcription across aging in C57Bl/6J mice. Neurobiol Aging 130, 141–153 (2023). 10.1016/j.neurobiolaging.2023.05.015

96 Gruene, T. M., Flick, K., Stefano, A., Shea, S. D. & Shansky, R. M. Sexually divergent expression of active and passive conditioned fear responses in rats. Elife 4 (2015). 10.7554/eLife.11352

97 Colom-Lapetina, J., Li, A. J., Pelegrina-Perez, T. C. & Shansky, R. M. Behavioral Diversity Across Classic Rodent Models Is Sex-Dependent. Front Behav Neurosci 13, 45 (2019). 10.3389/fnbeh.2019.00045

98 Borkar, C. D. et al. Sex differences in behavioral responses during a conditioned flight paradigm. Behav Brain Res 389, 112623 (2020). 10.1016/j.bbr.2020.112623

99 Anagnostaras, S. G. et al. Automated assessment of pavlovian conditioned freezing and shock reactivity in mice using the video freeze system. Front Behav Neurosci 4 (2010). 10.3389/fnbeh.2010.00158

100 Malliaros, F. D. & Vazirgiannis, M. Clustering and community detection in directed networks: A survey. Phys Rep 533, 95–142 (2013). 10.1016/j.physrep.2013.08.002

101 Rubinov, M. & Sporns, O. Complex network measures of brain connectivity: uses and interpretations. Neuroimage 52, 1059–1069 (2010). 10.1016/j.neuroimage.2009.10.003

102 Phillips, R. A., 3rd et al. An atlas of transcriptionally defined cell populations in the rat ventral tegmental area. Cell Rep 39, 110616 (2022). 10.1016/j.celrep.2022.110616

103 Young, M. D. & Behjati, S. SoupX removes ambient RNA contamination from droplet-based single-cell RNA sequencing data. Gigascience 9 (2020). 10.1093/gigascience/giaa151

104 Germain, P. L., Lun, A., Garcia Meixide, C., Macnair, W. & Robinson, M. D. Doublet identification in single-cell sequencing data using scDblFinder. F1000Res 10, 979 (2021). 10.12688/f1000research.73600.2

105 Hao, Y. et al. Dictionary learning for integrative, multimodal and scalable single-cell analysis. Nat Biotechnol 42, 293–304 (2024). 10.1038/s41587-023-01767-y

106 Yao, Z. et al. A high-resolution transcriptomic and spatial atlas of cell types in the whole mouse brain. Nature 624, 317–332 (2023). 10.1038/s41586-023-06812-z

107 Smith, M. S., Freeman, M. E. & Neill, J. D. The control of progesterone secretion during the estrous cycle and early pseudopregnancy in the rat: prolactin, gonadotropin and steroid levels associated with rescue of the corpus luteum of pseudopregnancy. Endocrinology 96, 219–226 (1975). 10.1210/endo-96-1-219

